# From flagellar motor behavior to bacterial swimming: defining a reference state for motility dynamics in *Magnetospirillum gryphiswaldense*

**DOI:** 10.64898/2026.09.09.750318

**Authors:** Diego Roesch, Gabriel Delabre, Émilie Gachon

## Abstract

Cell motility is powered by the bacterial flagellar motor, a rotary nanomachine whose activity is dynamically modulated by external stimuli. In *Magnetospirillum gryphiswaldense*, magnetic and chemical inputs are thought to converge at the level of motility control, yet the mechanisms underlying chemotactic regulation remain poorly understood. Here, we seek to establish a quantitative reference-state of a model strain of magnetotactic bacteria through bacterial flagellar motor dynamics. To achieve this goal, magnetotaxis was kept as a natural factor by keeping the earth’s magnetic field as the only source of magnetism. To avoid aerotaxis bias, oxygen gradients were removed by implementing two different constant oxygen conditions: environmental oxygen exposure, and limited oxygen exposure. Together, the tethered-cell bacterial flagellar rotational assay and the free-swimming assay in the absence of external stimuli presented in this paper establish a framework for investigating motor and cellular swimming adaptation to magnetic, aerotactic, and chemical signals. Under reference-state conditions, the bacterial flagellar motor showed a tendency to exhibit log-normal distributions for the time spent in each motor state: runs in different directions (counter-clockwise and clockwise), and pause. A semi-Markov reference-state model was developed to provide a quantitative description of bacterial flagellar motor dynamics. The model revealed that cell magnetic polarity modulates the transition pathways leading to the paused state, whereas oxygen, in the absence of a gradient, primarily regulates residence in each motility state.

## 1 Introduction

### 1.1 Magnetotactic bacteria as a model for multisensory navigation

Magnetotactic bacteria (MTB) comprise a diverse group of microorganisms distributed across multiple bacterial lineages [1, 6]. These bacteria have attracted considerable attention because of their remarkable ability to use environmental iron to biomineralize intracellular magnetic nanoparticles [11, 33, 40]. These nanoparticles are commonly composed of greigite (*Fe*_3_*S*_4_) or magnetite (*Fe*_3_*O*_4_) and are enclosed within lipid-bilayer vesicles called magnetosomes, which are typically arranged in linear chains along the longitudinal axis of the cell [14, 36, 40]. The magnetosome chain gives the cell a permanent magnetic dipole, allowing the classification as north-seeking (NS) when they swim towards the magnetic north, and south-seeking (SS), when they swim towards the magnetic south [35, 3].

Bacterial motility is strongly influenced by the chemical and physical characteristics of the natural environment. The ability of bacteria to sense their surroundings is essential for survival and is therefore central to understanding their motility patterns. Bacterial trajectories are shaped by different forms of taxis, defined as directed behavioral responses that bias cell movement toward or away from specific environmental stimuli. Among these, chemotaxis enables bacteria to navigate chemical gradients by increasing their residence in favorable regions containing attractants, such as nutrients, and avoiding regions containing repellents [41, 49]. MTB commonly inhabit chemically stratified aquatic environments near the oxic-anoxic transition zone (OATZ) and therefore exhibit aerotaxis, a specialized form of chemotaxis that enables them to navigate oxygen gradients and locate their preferred microaerobic habitat [34, 11, 12, 23]. In addition to responding to oxygen, redox taxis enables MTB to bias their movement toward metabolically favorable oxidation-reduction conditions, helping them locate and remain within suitable zones along the steep redox gradients characteristic of stratified aquatic environments [22]. MTB also possess the remarkable ability to align with the Earth’s magnetic field through magnetotaxis. Unlike chemotaxis, or redox taxis, magnetotaxis does not involve attraction to or repulsion from magnetic fields meaning it is not a real taxis; rather, the magnetic field provides a directional cue that constrains the bacteria’s cell body along magnetic field lines thereby facilitates the search for favorable chemical and oxygen conditions [12, 11, 47]. These mechanisms are not mutually exclusive, and MTB integrate multiple taxis and magnetic alignment to navigate efficiently through natural habitats characterized by complex geochemical gradients and physical barriers [22, 25, 23].

In MTB, a central biological question is how magnetic orientation and chemically derived signals are integrated to produce a coordinated motility response. Magnetotactic polarity is not necessarily fixed, as changes in oxygen availability or other chemical conditions can alter the preferred swimming direction and induce reversals relative to the magnetic field [31, 25, 23]. In MSR-1, matching population swimming polarity to the field orientation improves migration into an aerotactic band, providing direct evidence that magnetic alignment and oxygen sensing function together during navigation [42]. However, the cellular mechanisms through which these different inputs are coordinated remain poorly understood. Addressing this question requires investigating the machinery that converts environmental information into changes in motility, particularly the chemotaxis signaling network and the bacterial flagellar motor (BFM).

### 1.2 The bacterial flagellar motor as the output of sensory integration

Within the canonical diderm bacterial model, the BFM is organized around two main functional modules: the rotor and the stator units. The rotor includes the MS ring embedded in the cytoplasmic membrane, the C-ring on the cytoplasmic side, the rod, and the P and L rings that stabilize the rotating shaft across the peptidoglycan layer and outer membrane [18, 48, 16]. The C ring is particularly important because it participates in torque transmission, directional switching, and chemotaxis-dependent regulation [17, 37]. Stator units, arranged around the rotor in the cytoplasmic membrane, conduct ions such as H^+^ or Na^+^ and convert ion flow into mechanical torque; in MSR-1, this process is driven by H^+^ ions [48].

Chemical cues trigger signaling pathways in MTB that appears to follow the canonical *Escherichia coli* K-12 RP437 (*E*.*coli*) chemotaxis cascade, where chemotaxis-related proteins, commonly referred to as Che proteins and encoded by *che* genes, mediate signal transduction from chemoreceptors to the BFM [31, 47, 41, 8]. At the cell poles, chemoreceptors known as methyl-accepting chemotaxis proteins (MCPs) cluster into arrays and act as sensors that detect chemical cues. Ligand binding modulates the chemotaxis cascade by regulating the phosphorylation state of the histidine kinase CheA, which subsequently transfers the phosphoryl group to the response regulator CheY. Phosphorylated CheY (CheY-P) then diffuses to the BFM, where it binds to the C-ring and promotes switching between counterclockwise (CCW) and clockwise (CW) rotation, thereby modulating swimming behavior [4, 47]. In motile swimming bacteria, BFM rotation represents the output of the chemotaxis signaling cascade. It serves as a reliable proxy for chemical sensing and provides an experimentally tractable readout for elucidating how MTB integrate magnetic and chemical cues into a coordinated motility response.

### 1.3 Establishing the baseline dynamics of the MSR-1 flagellar motor

In many motile bacteria, navigation emerges from the alternation between periods of directed swimming and stochastic reorientation events. The canonical example is the run-and-tumble behavior of *E*.*coli* [2], although many species, including MTB, employ distinct swimming and reorientation strategies [49, 7, 44, 32, 29]. In *E*.*coli*, counterclockwise (CCW) rotation promotes the formation of a flagellar bundle that propels the cell during a run, whereas clockwise (CW) rotation of one or more motors disrupts the bundle and produces a tumble resulting in a stochastic reorientation of the cell body. [7, 39]. However, this precise relationship between motor rotation and whole-cell movement is not universal. For example, in *Vibrio alginolyticus*, CW and CCW rotation of the polar flagellum are associated with forward and backward swimming, respectively [24]. In MSR-1, to understand the relationship between motor rotation and swimming behavior it is necessary to take into consideration multiple factors including the helical morphology of the cell body, the coordination of the cell’s bipolar flagella, and the passive magnetic alignment of the cell body to magnetic field lines. Consequently, the motor states and switching patterns of MSR-1 cannot be inferred directly from the motility models established for non-magnetic bacteria.

Before determining how magnetic and chemical cues modify MSR-1 motility, it is therefore necessary to establish the reference-state dynamics of its BFM. A quantitative characterization of these dynamics is currently lacking, making it difficult to distinguish stimulus-induced responses from the intrinsic variability and temporal organization of the motor. Several complementary measurements are required to define this baseline. Rotational speed quantifies the magnitude of motor output, while switching dynamics describe the frequency and temporal organization of transitions between CW, CCW, and pause states [38]. Rotational bias, defined by the relative occupancy of CW and CCW rotation, identifies whether the motor preferentially operates in one direction [38]. Pause frequency and duration further characterize transient interruptions in rotation of one of the motors that may contribute to changes in swimming behavior such as direction and speed. Together, these measurements define the normal operating regime of the BFM and provide a quantitative reference against which responses to controlled perturbations can be evaluated.

Accordingly, the objective of this work is to establish a quantitative reference state for the MSR-1 BFM under defined oxygen exposure and constant magnetic-field conditions, in the absence of an imposed chemical stimulus. By characterizing switching dynamics, directional bias, and pause behavior, this baseline will provide the framework needed to determine how future magnetic and chemical perturbations reshape motor-level outputs. Establishing this reference state is therefore a necessary first step toward understanding how multiple environmental cues are coordinated through the chemotaxis signaling network and expressed as changes in the BFM behavior.

### 2 Materials and Methods

### 2.1 *Magnetospirillum gryphiswaldens* culturing

MSR-1 was cultured in Flask Standard Medium (FSM) prepared in-house. All chemicals were acquired from Sigma-Aldrich. FSM contained HEPES buffer 10 mM, lactate, soy peptone, yeast extract, NaC_3_H_5_O_3_, KH_2_PO_4_, MgSO_4_, C_6_H_5_FeO_7_, and Wolfe’s mineral solution. The medium was sterilized by autoclaving at 121.1°C for 15 minutes, and the pH was adjusted to 7.0 with sterile NaOH. Cultures were prepared by adding 10mL of FSM to sterile 15mL Hungate tubes, followed by inoculation with MSR-1 strain R3/S1 from a frozen stock. Tubes were fitted with a needle connected to a 0.22 µm filter through the septum to allow gas exchange with the headspace. Cultures were incubated statically at 28°C for 24 hrs and used during exponential growth, corresponding to an optical density of 0.08–0.20 at 600nm.

For the experiments, 1 mL of the culture was sheared using two syringes connected through a silicone tubing with internal diameter of 0.5 mm and a needle gauge 23, the MSR-1 solution was pushed between syringers 25 times, followed by centrifugation at 5000g for 5 minutes to pellet the cells. The supernatant was discarded, and the pellet was resuspended in 1 mL of motility buffer which consiste of 10 µM HEPES buffer at pH 7.0 ± 0.2.

The same culturing protocol was followed for tethered experiments and for the free swimming experiments, but for the latter, the bacteria was not sheared, only resuspended in HEPES buffer 10 mM after centrifugation.

### 2.2 Tethering of bacteria

Tethering was performed using poly-L-Lysine (PLL) 0.1% w/v (150KDa - 300KDa) from Sigma-Aldrich as ionic agent to fixate the flagella from MSR-1 to the surface of a coverslip. The coverslip was immersed in 0.005% (v/v) PLL solution for 20 minutes to ensure complete coating, and subsequently rinsed thoroughly with deionized water. To properly tether by the flagella, the coverslip was placed on top of the north or south pole of a magnet and 10 µL of MSR-1 in motility buffer was placed in the center of the coverslip for 20 minutes in a closed chamber. The incubation along the magnet allowed proper “Z” axis alignment of MSR-1 promoting contact between the flagella and the surface as depicted in Figure 1.

**Figure 1.**
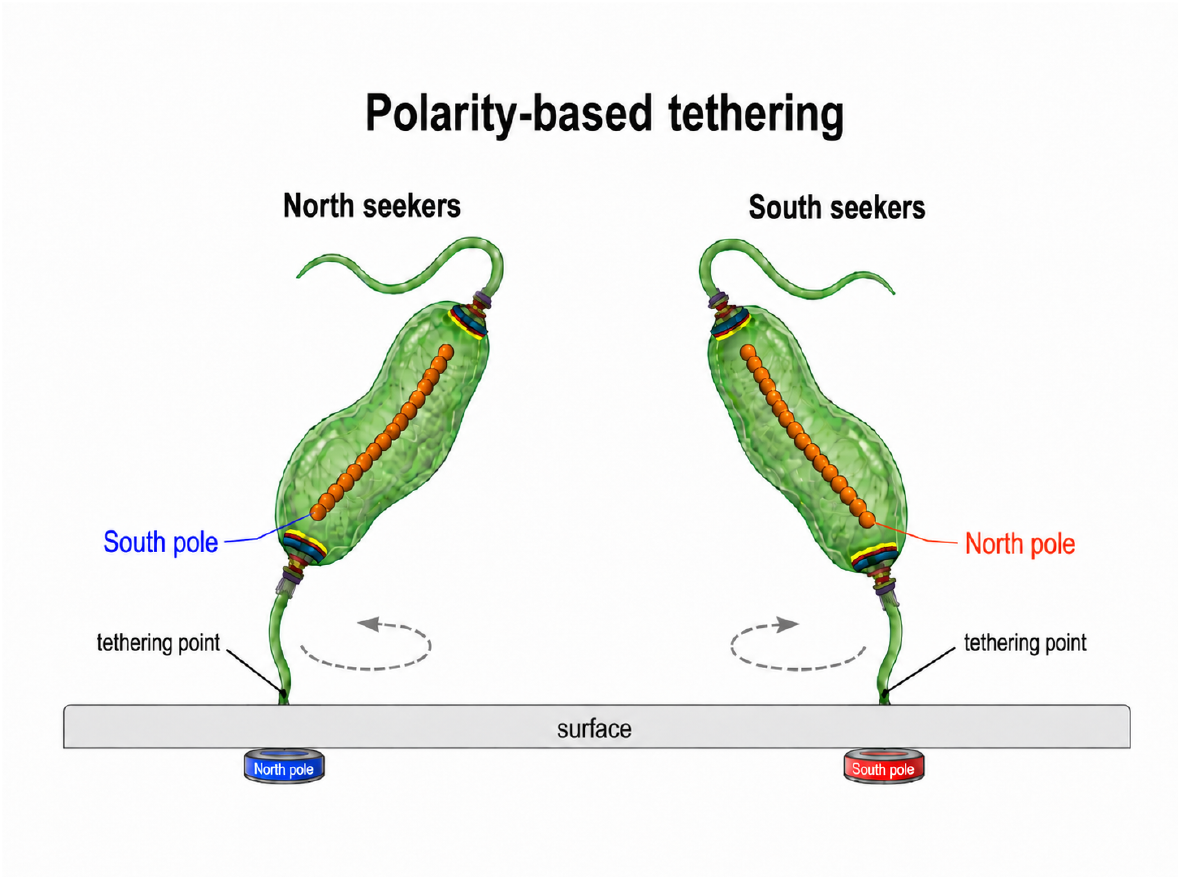
Schematic scheme for specific tethering of north-seeking and south-seeking cells. The magnets allow for passive allignment of the cell via the magnetosome chain, favoring the flagella interation to with the surface for the specific cell polarity.

The bacteria was enclosed inside a circular chamber (13.9 mm diameter and 200 µm height) made of a polyethylene terephthalate (PET) double sided tape between a microscope slide and the coverslip with the tethered bacteria.For the experiments two configurations were followed:

1. The chamber was opened by cutting the tape on one side, allowing constant exposure to environmental air.
2. The chamber was kept intact to create a closed environment with limited oxygen.

The first configuration allowed for a constant oxygen concentration which was referred as Environmental O_2_ condition, while the second approach is referred to as Low O_2_ condition due to the limited oxygen availability and constant oxygen consumption of the culture.

The previous scheme was reproduced for both north seekers and south seekers, by changing the magnet orientation. Therefore four different configurations were created: north and south seekers under environmental O_2_ condition and Low O_2_ condition.

### 2.3 3D swimming experiments

For the 3D swimming experiments, MSR-1 culture was subjected to magnetic fields from fixed magnets allowing the separation betweeen north seekers and south seekers. Only south seekeres where selected for the experiments. To study free swimming both conditions Low O_2_ (2%), and Environmental O_2_ (21%) were studied. For environmental O_2_ the double sided tape scheme was reproduced using an oxygen permeable silicone double sided tape with a circula chamber of 16 mm diameter and 210 µm height. For low O_2_ condition, a polydimethylsiloxane (PDMS) microfluidic chips was used. The chip consisted of 3 channels, where the side channels were flushed with a constant mixture of nitrogen and oxygen for a final O_2_ concentration of 2%. The chip was coated entirely with the exception of the visualization area with a vinylpolysiloxane resin to avoid oxygen permeability from the environment, and the visualization area had an additional coverslip inserted on top to avoid oxygen difussion from the environment promoting a constant 2% O_2_ saturated environment through the chip [43].

### 2.4 Microscopy & 2D Video Tracking

Cell rotation was recorded in bright-field microscopy using an in house build inverted microscope with a 40x objective with water immersion and a high-speed camera Kinetix (1T-01-N-KINETIX-MC) at 250 frames per second (fps) over 5-minute intervals.

### 2.5 Microscopy & 3D Video Tracking

3D swimming trajectories were recorded at the ESPCI-PMMH Sorbornne University with a modified Zeiss Observer Z1 inverted microscope coupled to a 3D Lagrangian tracking platform, allowing for automatic movement in the X,Y,Z planes to follow swimming bacteria at the single-cell level [5, 43]. The microscope was equipped with a 63x water-immersion objective, an adaptative sample holder for microfluidic chips and a high-speed camera (Hamamatsu Orca-Flash 4.0 CMOS) operating at 80 fps. The system was configured to capture the three-dimensional movement of MSR-1 cells in real time, allowing for detailed analysis of their swimming behavior under specific oxygen conditions. Each track lasted as long as the bacteria didn’t escaped field of view.

### 2.6 Data Analysis

#### 2.6.1 Tethered bacteria data analysis

Rotational analysis of MSR-1 was carried out using BRAS software. The software determines position coordinates for each frame by computing the angular displacement of the cell’s center of mass [20].

Singel-cell rotational trajectories were calculated from coordinates in the HD5 files generated by the BRAS software. The raw center of mass coordinates were corrected for experimental drift and geometric distortion before estimating the angular displacement. For each trajectory, local drift was corrected by fitting the data into ellipses and substracting the estimated local center. The drift corrected trajectory was then aligned to the best ellipse, and subjected to a whitening transformation using the covariance structure of the trajectory, and soft normalization of the radius to fit into a circular trajectory. The corrected coordinates were converted into angular positions using the four quadrant arctangent function. The obtained angular position was then unwrapped to obtain a continuous angular trajectory, and the instantaneous angular speed was calculated as the discrete derivative of the unwrapped angular position.

The unwrapped angular signal was expressed as cumulative turns:

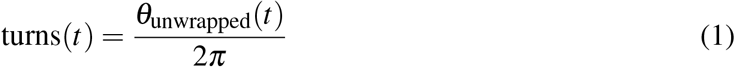

where *θ*_unwrapped_ is the corrected angular position in radians. A rough instantaneous rotational speed was also calculated from the frame-to-frame angular increment:

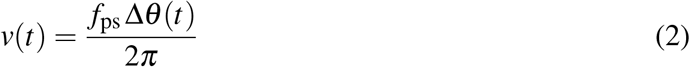

Motor dynamics were segmented from the cumulative turns signal instead of the instantaneous speed to avoid noise amplification from the differentiation step. The segmentation process was performed through the Pruned Exact Linear time Change-Point algorithm (PELT) with a L2 Cost model, and a penalty of 7 [19]. A piecewise linear fit was applied to each segment between change points to acquire the slope for each segment. All the segments were classified unsing a symmetric pause band around zero speed, where segments with fitted speed above +0.25 Hz were assigned to clockwise rotation, frames below −0.25 Hz were assigned to counterclockwise rotation, and frames within ±0.25 Hz were assigned to a pause state. In the current implementation, positive slopes are labeled CW and negative slopes are labeled CCW.

After segmentation, each continuous interval classified as CW rotation, CCW rotation, or pause was treated as an individual motor-state event. For each event, the state identity, duration, fitted rotational speed, chronological position, cell identifier, magnetic polarity, and oxygen condition were retained. The analysis was conducted at two complementary levels: a cell level, in which every cell contributed one summary value for each metric, and an event-pooled level, in which all events from cells belonging to the same experimental condition were combined.

Pause events were further classified according to the motor-rotation direction immediately before and after the pause. Pauses followed by rotation in the direction opposite to that preceding the pause were classified as direction-switching pauses (CW–pause–CCW or CCW–pause–CW). Pauses followed by rotation in the same direction were classified as direction-retaining pauses (CW–pause–CW or CCW–pause–CCW).

At the cell level, the number of CW runs, CCW runs, direction-retaining pauses, and direction-switching pauses was calculated separately for every cell. The mean and median duration of each event class were also calculated within each cell. Condition-level summaries were subsequently calculated across cells and included the mean, median, standard deviation, first and third quartiles, and nonparametric bootstrap confidence intervals. Because the duration metrics were generally right-skewed, the median was used as the principal measure of central tendency for the cell-level duration comparisons.

For the event-pooled analyses, all event durations from cells belonging to the same combination of magnetic polarity and oxygen condition were combined. These pooled events were used to characterize the condition-level dwell-time and inter-switch-interval distributions. Candidate probability distributions were fitted to the pooled positive event durations by maximum likelihood, and their agreement with the observed data was evaluated using the Kolmogorov–Smirnov (KS) statistic and Bayesian information criterion (BIC), as reported in Table S1. The lognormal model provided the best-supported description for the principal dwell-time distributions. For a lognormal random variable, the probability density was defined according to Equation 3, where *x* is the event duration in seconds, *µ* is the mean of ln(*x*), and *σ* is the standard deviation of ln(*x*). The fitted distribution parameters are reported in Table S2. Because these distribution fits pool all events within each condition, they provide an event-weighted description of the recorded population.

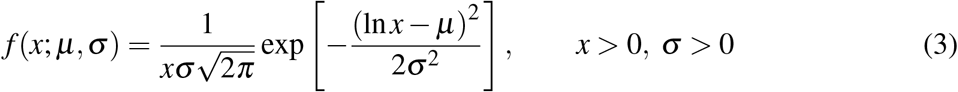

The final reference-state model of BFM rotational dynamics in MSR-1 was constructed by integrating the statistical properties of each rotational state, including pause subclasses defined by their associated switching dynamics. Since the dwell-time distributions of the observed events were right-skewed and better described by lognormal-like distributions than by memoryless exponential distributions, a semi-Markov model was selected instead of a conventional Markov chain [30]. The model combined an embedded transition probability matrix, describing transitions among rotational states, with state-specific dwell-time distributions, describing how long the motor persisted in each state. Long-term state occupancy was estimated by weighting the stationary probabilities of the embedded chain by the corresponding mean dwell times. This framework therefore captured both the transition structure and the non-memoryless residence-time behavior of the BFM under reference-state conditions.

#### 2.6.2 Swimming bacteria data analysis

Three-dimensional single-cell trajectories were obtained using an AI-assisted Lagrangian tracking microscope adapted for bright-field imaging of MSR-1. Real-time bright-field images were processed by a convolutional neural network trained on controlled Z-scan image libraries of inactive MSR-1 cells, acquired from −15 to +15 µm around the focal plane [43]. The network estimated the instantaneous displacement of the bacterium from the centered and focused reference position, returning Δ*X*, Δ*Y*, and Δ*Z* corrections. These corrections were sent to a LabVIEW feedback routine, synchronized with image acquisition by a TTL trigger, which commanded the mechanical X,Y stage and piezoelectric Z actuator to recenter and refocus the bacterium. Because the bacterium was kept approximately fixed in the camera frame by the feedback loop, its laboratory-frame trajectory was reconstructed from the time series of absolute stage positions recorded during tracking. The resulting data consist of X(t), Y(t), and Z(t) coordinates of individual MSR-1 cells, together with the corresponding image sequence, enabling subsequent calculation of displacement, velocity, persistence, and reversal events [43].

The raw X,Y, Z coordinates were obtained from the lagrangian AI tracking system as mentioned above. The raw coordiantes were smoothed using a fixed one dimensional Gaussian filter with *σ* = 1.5 to reduce high-frequency noise.The smoothed trajectories were centered by substracting the average position from each point, and projected onto its dominant axis of motion using the principal components analaysis (PCA). The one-dimensional projection was further smoothed with a Gaussian filter with *σ* = 2.0 frames.

The Bacterial trajectories for 3D swimming were analyzed using a custom python pipeline. The length of each track varied depending on how long the bacterium could be kept within the chamber, therefore to avoid bias from track lenght, all the tracks where segmented into 25 second windows. The metrics were calculated based on hierarchy calculating all statistics for each window, followed by a statistical analysis of all the windows per track to yield aggregate statistics per sample. Each trajectory was converted into a chronological sequence of observed motility events. Pauses and direcitonal reversals were identified by speed reductions below the 5th percentile of the track-specific speed distribution, representing a drastic speed reduction relative to the typical swimming speed of that track. Directional reversals were identified as local extrema in the smoothed PCA projection by detecting both peaks and troughs using a prominence criterion. The prominence threshold was set to twice the median absolute frame-to-frame change in the smoothed projection. Consecutive low-speed frames were classified as pauses. Non-pause intervals were classified as forward or backward runs according to the sign of displacement along the PCA projection axis. Adjacent events with the same state label were merged, producing a compact observed-state sequence composed of three states: Forward, Pause, and Backward. For each event, the pipeline recorded start time, end time, duration, speed, and event type.

The analysis of the processed trajectories allowed for the determinations of multiple metrics included mean and median speed, reversal frequency, pause frequency, reversal count, pause count, run count, forward-event count, backward-event count, pause-event count, total event count. State-transition dynamics were analyzed using both embedded Markov and observed-state semi-Markov summaries. The chronological event sequence was first compressed into a transition sequence over the states Forward, Pause, and Backward. Transition counts were calculated between consecutive events and row-normalized to generate embedded transition probability matrices at per-sample and pooled levels. These matrices describe the order of state transitions independently of state residence times.

To determine whether a regular Markov interpretation was appropriate, dwell-time distributions were analyzed for each observed state. For Forward, Pause, and Backward events, dwell durations were fitted by maximum likelihood to exponential, gamma, lognormal, and Weibull distributions. Models were compared using the BIC and Kolmogorov–Smirnov goodness-of-fit statistics as reported in Table S3. A regular Markov interpretation was accepted only when all three state dwell-time distributions were best described by the exponential model according to BIC and when the exponential Kolmogorov–Smirnov test was not rejected at *p <* 0.05. If any state failed this criterion, the process was classified as a semi-Markov process, retaining the embedded transition matrix as the transition skeleton while explicitly reporting state-specific dwell-time statistics.

For condition-level summaries of the free-swimming data, the 25-second windows were first aggregated within each trajectory so that every tracked cell contributed one value for each metric. The distributions of these per-cell metrics were assessed using graphical inspection together with the Shapiro–Wilk and Anderson–Darling tests. Approximately symmetric metrics were summarized using the mean and standard deviation, whereas right-skewed metrics were summarized using the median and interquartile range. For median-based summaries, 95% confidence intervals were calculated using 3,000 nonparametric bootstrap resamples of complete cell trajectories. Windows from the same trajectory were not treated as independent biological replicates.

## 3 Results

### 3.1 Baseline of the bacterial flagellar motor dynamics exhibit non-Markovian temporal behavior

To characterize baseline motor dynamics, we analyzed the rotational behavior of the BFM in MSR-1 under reference-state conditions. These conditions were defined by the absence of externally added chemical stimuli, controlled oxygen availability, and exposure to the ambient Earth’s magnetic field. The analysis focused on state occupancy, state-transition sequences, pause-duration distributions, and inter-switch interval distributions.

Across the analyzed conditions, both pause durations and time between switching events exhibited right-skewed distributions, characterized by frequent short states and a smaller number of long-duration states. In Figures 2 and 3, the lognormal model provided the best description for the experimental distributions, as supported by the quantitative result in Table S1. Across all tested conditions, the lognormal model consistently produced the lowest Kolmogorov-Smirnov (KS) statistic, and lowest Bayesian information criterion (BIC), indicating that both types of event durations were better described by right-skewed, multiplicative-like distributions than by normal, exponential, Weibull, or gamma models.

**Figure 2.**
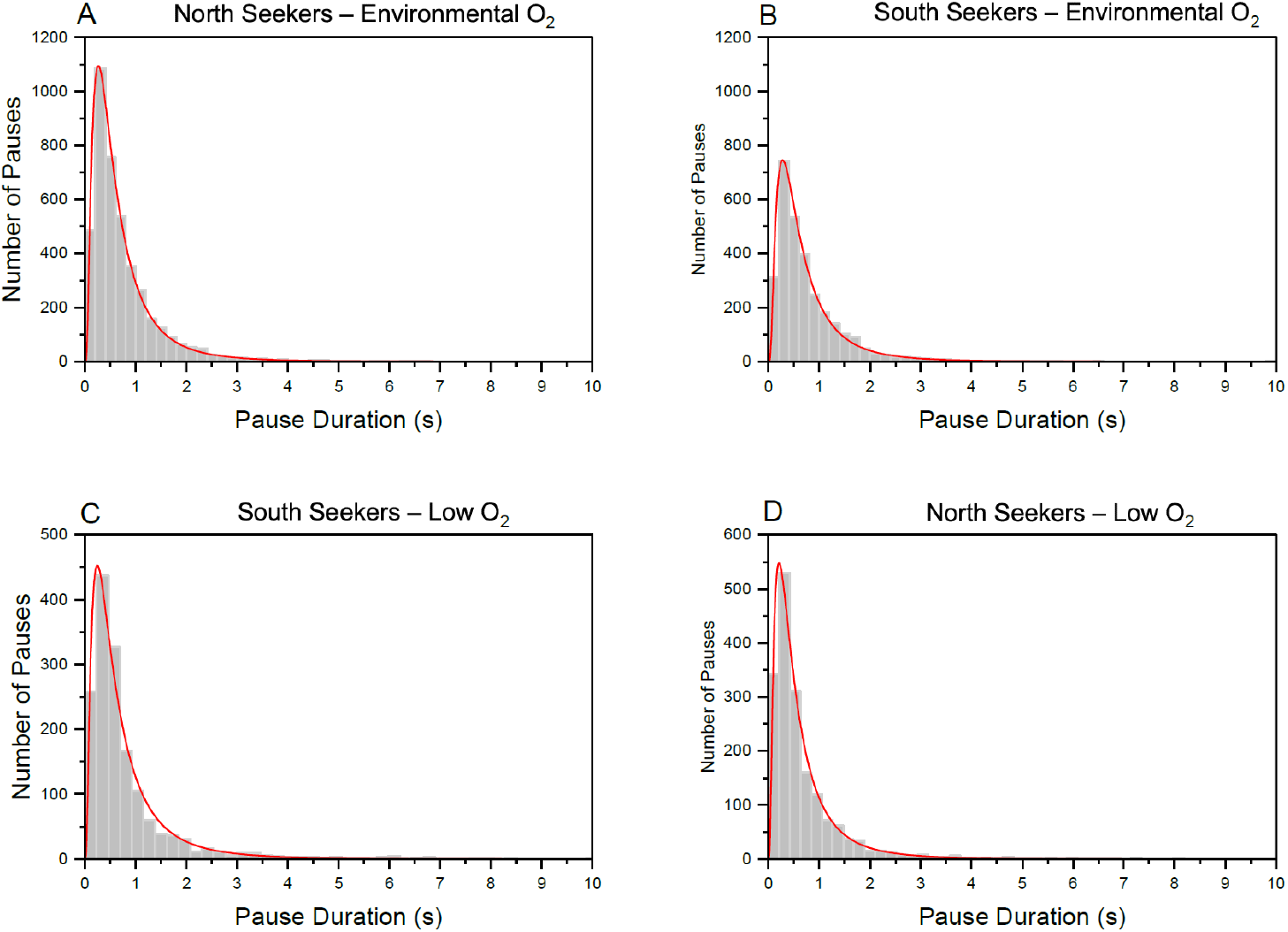
Lognormal distribution fit for pause durations under low O_2_ and environmental O_2_ conditions. The fit parameters and goodness-of-fit statistics are reported in Table S1 and S2. (A) Probability density function of pause durations for north seekers under environmental O_2_ condition. (B) Probability density function of pause durations for south seekers under environmental O_2_ condition. (C) Probability density function of pause durations for north seekers under low O_2_ condition. (D) Probability density function of pause durations for south seekers under low O_2_ condition.

**Figure 3.**
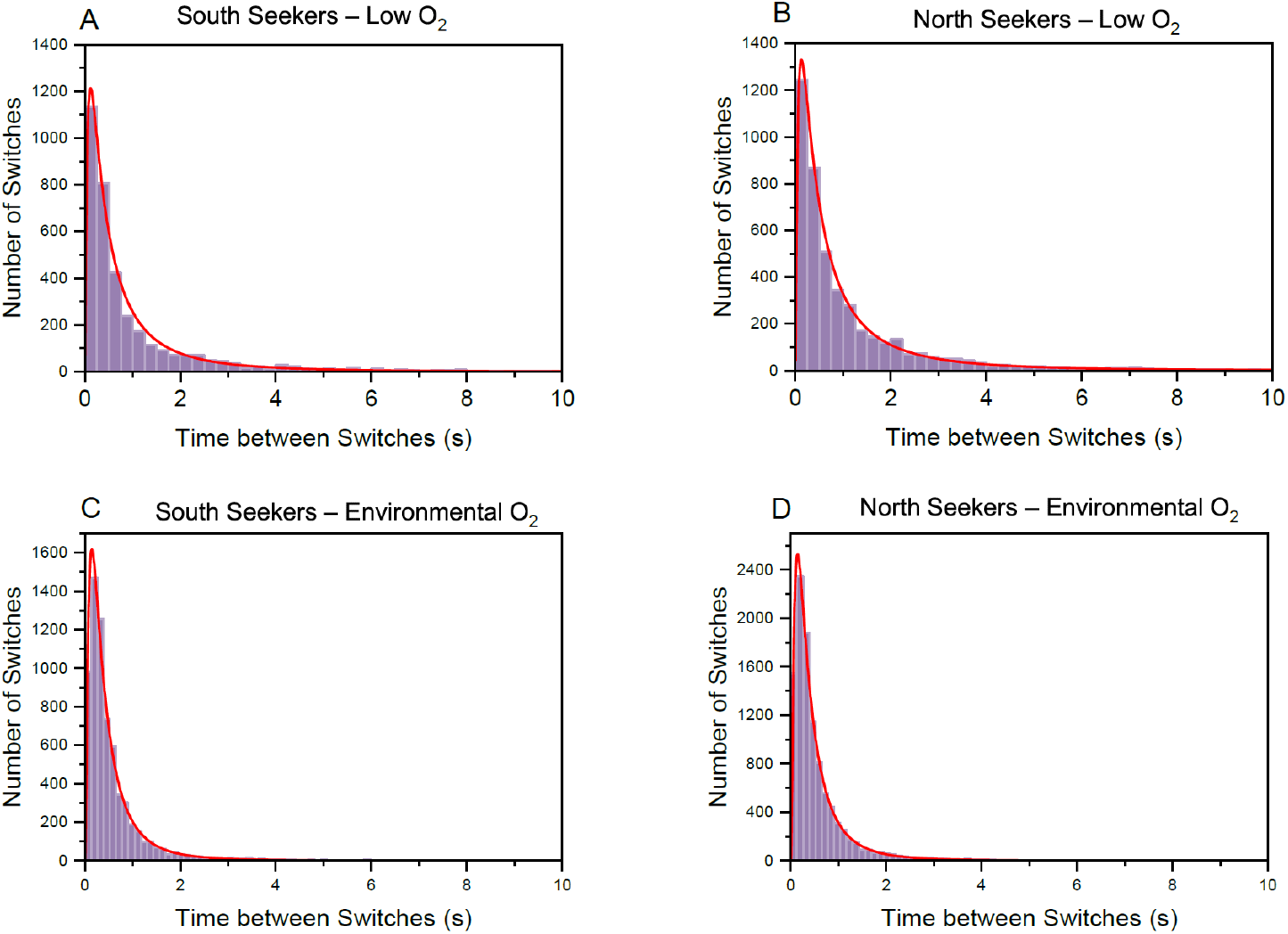
Lognormal distribution fit for switch durations under low O_2_ and environmental O_2_ conditions. The fit parameters and goodness-of-fit statistics are reported in Table S1 and S2. (A) Probability density function of switch durations for north seekers under environmental O_2_ condition. (B) Probability density function of switch durations for south seekers under environmental O_2_ condition. (C) Probability density function of switch durations for north seekers under low O_2_ condition. (D) Probability density function of switch durations for south seekers under low O_2_ condition.

Direct transitions between CCW and CW rotation accounted for the largest fraction of transition states, whereas transitions that occur after a pause (pause-mediated) occurred at lower but consistent frequencies across conditions, as shown in Figure 4. Overall, the transition patterns in Figure 3A were broadly similar between cell polarities (north and south leading poles), and across oxygen conditions. However, in Figure 4B the state occupancy analysis revealed condition-dependent differences in directional bias. Under low-O_2_ conditions, NS cells displayed higher CW occupancy than SS cells, whereas under environmental O_2_ conditions, SS cells showed higher CW occupancy than NS cells. When comparing oxygen conditions, low O_2_ was associated with a 2–3% reduction in pause-mediated transitions and an approximately 15% increase in CW directional bias, suggestings that CW rotation represent the run state in the frontal motor.

**Figure 4.**
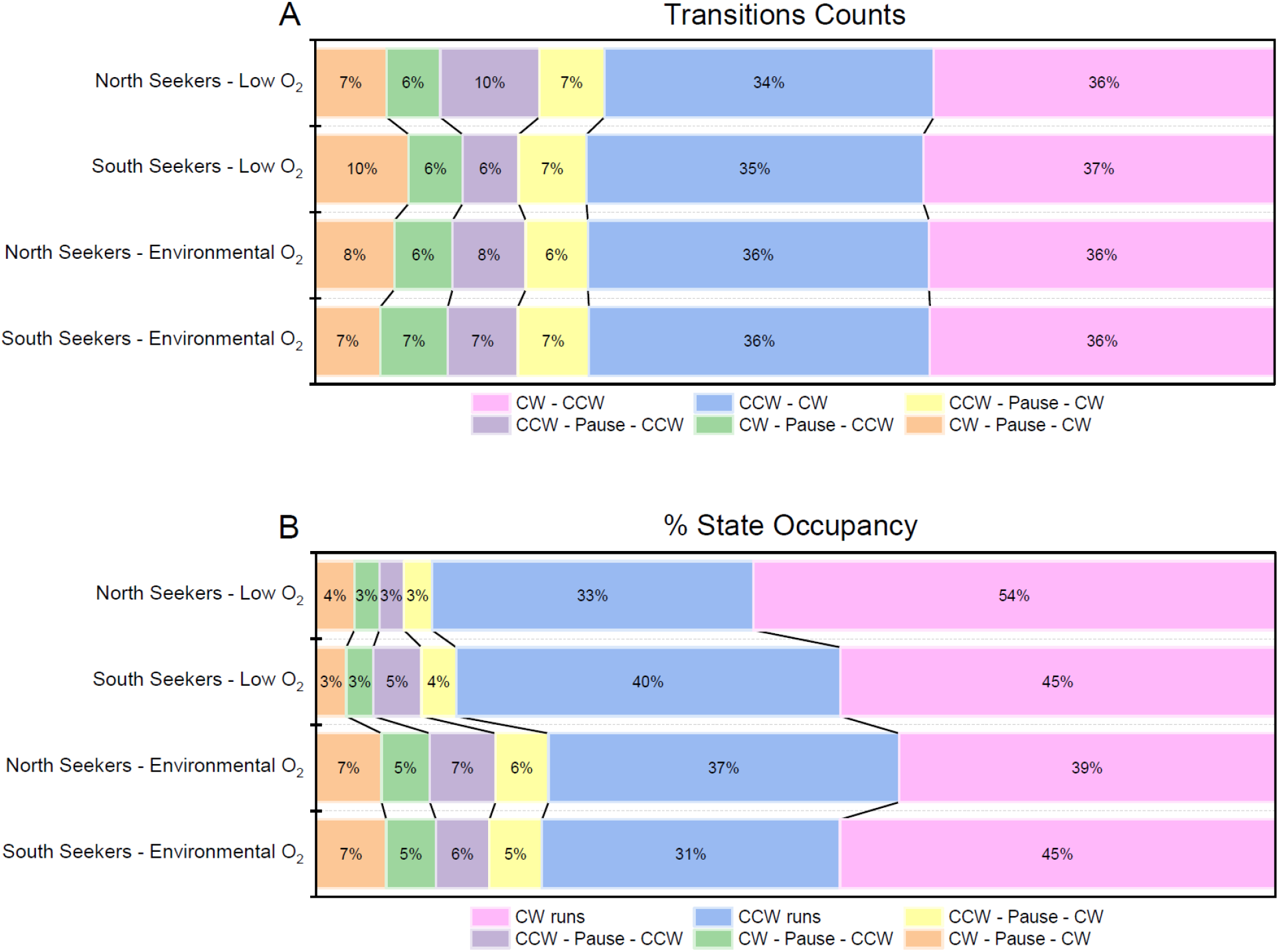
Transition counts and state occupancy of MSR-1 bacterial flagellar motor dynamics under environmental O_2_ and low O_2_ conditions. (A) Relative transition counts between CW runs, CCW runs, and pause-mediated transition states in North-seeking and South-seeking cells (B) Percentage state occupancy across the same conditions, showing the time spent in CW runs, CCW runs, and pause-associated states.

Pause and run dynamics were analyzed by separating state frequency from state occupancy, allowing us to distinguish changes in the rate of entering each motility state from changes in the duration of time spent in that state. Pauses were classified according to their effect on swimming direction: direction-switching pauses, in which the motor resumed rotation in the opposite direction (CW–pause–CCW or CCW–pause–CW), and direction-retaining pauses, in which the motor resumed rotation in the same direction as before the pause. Figure 5B outlines that direction-switching pauses represented the majority of pause ocurrences across all conditions, accounting for an average of 56% of pause counts, whereas direction-retaining pauses occurrence was higher with an average of 54%. However, Figure 5A shows that the contribution of each type of pause to total pause duration was mainly influenced by the oxygen conditions and cell polarity. Under environmental O_2_, direction-switching and direction-retaining pauses contributed nearly equally to the total pause duration in both cell polarities. Under low-O_2_ conditions, this balance became more seeker-dependent, with SS showing a greater duration contribution from direction-switching pauses, whereas NS showed a greater contribution from direction-retaining pauses.

**Figure 5.**
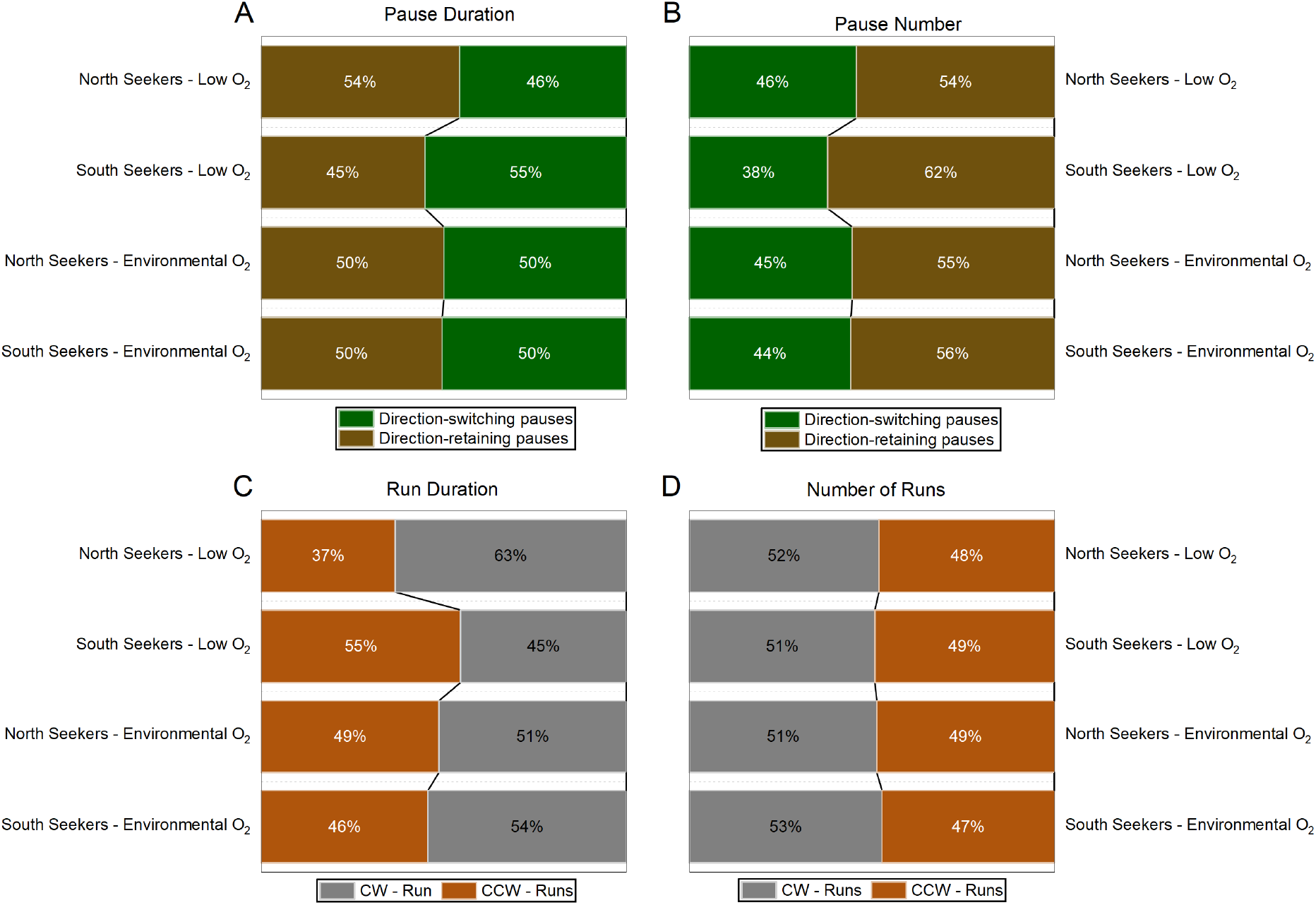
Cell level summaries of pause and run dynamics for north and south seekers under exposure to different oxygen conditions. Each parameter was based on the median value of cells per type of event. (A) Pairwise-normalized median pause duration based on the pause directional mechanism: direction-switching and direction-retaining. (B) Pairwise-normalized median number of pause events according to the pause directional mechanism: direction-switching or direction-retaining. (C) Pairwise-normalized median run duration based on the run directional mechanism: Clockwise and Counter Clockwise. (D) Pairwise-normalized median number of run events according to the run directional mechanism: Clockwise and Counter Clockwise.

A similar distinction was observed for run events. Figure 5D highlights that the number of CW and CCW runs remained close to balanced across all experimental groups, indicating that both rotational directions occurred with comparable frequency. In contrast, the run-duration analysis in Figure 5C reveals more pronounced direction-dependent differences based on cell polarity for both oxygen conditions. Under low O_2_, NS cells showed a higher contribution from CW runs, whereas SS cells showed a higher contribution from CCW runs, and for environmental O_2_ conditions, the differences were less pronounced for NS cells but more pronounced for SS cells.

### 3.2 The semi-Markov model reveals oxygen and polarity-dependent motor dynamics

To model the observed motor-state dynamics, we used a semi-Markov chain framework that incorporated both the sequence of state transitions and the residence time associated with each state on a population level. This approach accounts for the observed lognormal dwell-time distributions, consistent with state residence times that violate the memoryless assumption of a Markov process. The model was therefore used to distinguish two complementary properties of the dynamics: the probability of moving from one state to another, represented by the arrows, and the expected fraction of time spent in each state, represented by the node occupancies as depicted in Figure 6.

**Figure 6.**
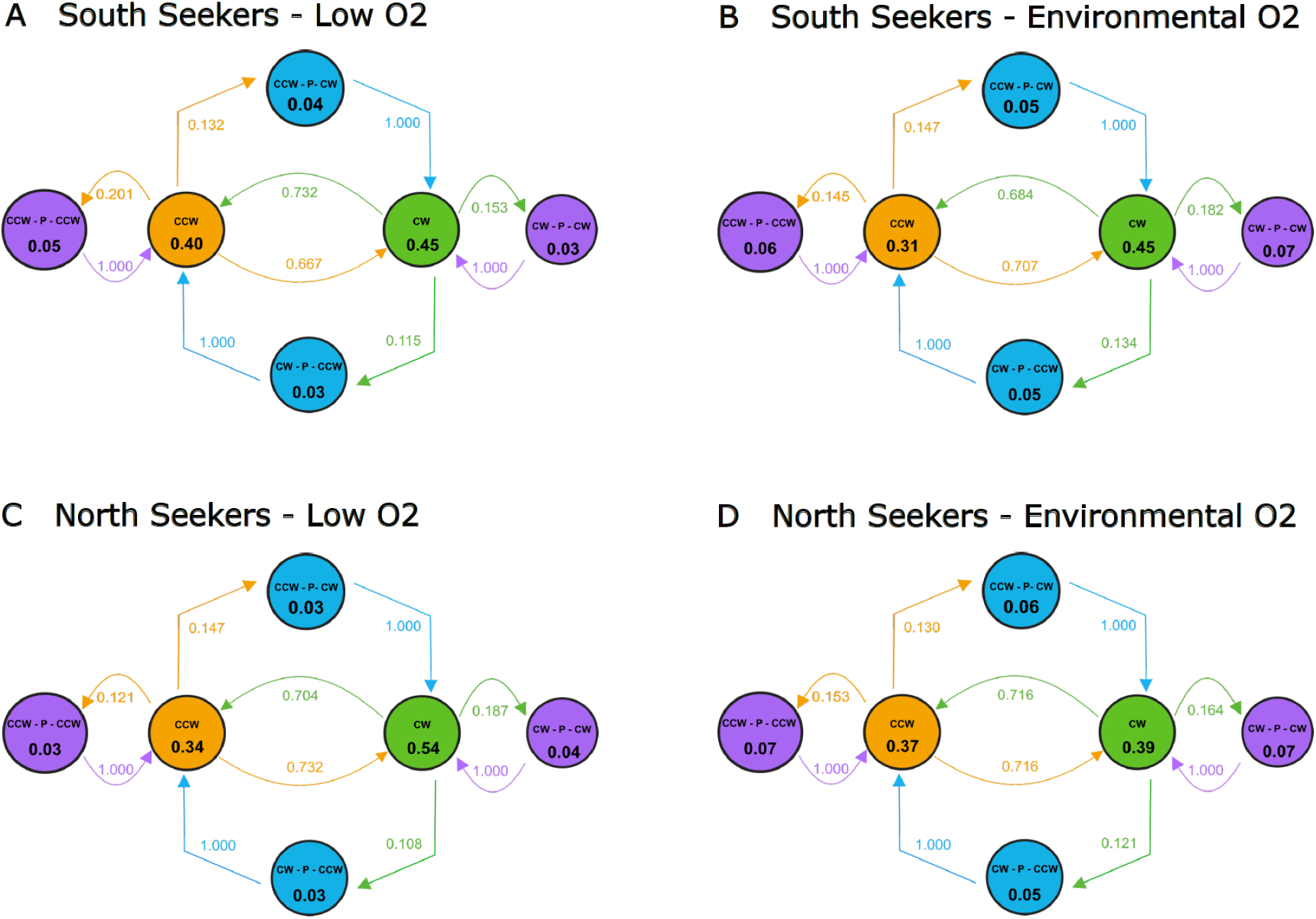
Semi-Markov model of the bacterial flagellar motor rotational dynamics in MSR-1 under steady-state conditions. All models consider the three main states: CW, CCW, and pause, and the subpause states: direction retaining pauses (CCW-P-CCW and CW-P-CW), and direction-switching pauses (CCW-P-CW and CW-P-CCW). (A) South seekers and (C) North seekers under low O_2_ conditions. (B) South seekers and (D) North seekers under environmental O_2_ conditions. The transition probabilities are represented by the arrows towards all the possible states. Complementing the transition probabilites, the dwell-time occupancy between all states over time is shown on each node.

Across all conditions, the semi-Markov chains showed that direct transitions between CCW and CW runs were the dominant switching route. This indicates that, under reference-state conditions, directional switching was primarily achieved through direct reversals between motor-rotation states, whereas pauses represented less frequent intermediate states. The probability of direct CCW-to-CW and CW-to-CCW transitions was consistently high, ranging from approximately 67% to 73%. In contrast, transitions from run states into pause-mediated pathways occurred with lower probabilities, generally between 11% and 20%. Because each pause node was defined by its entry and exit context, transitions out of pause states were deterministic in the diagram and should be interpreted as the completion of the corresponding pause-mediated pathway rather than as independent switching choices.

State occupancy analysis showed that oxygen availability altered motor-state residence in a polarity-dependent manner. Under low O_2_, both NS and SS cells favored the CW state, although this bias was markedly stronger in NS cells (CW: 54%, CCW: 34%) than in SS cells (CW: 45%, CCW: 40%). Under environmental O_2_, the CW bias persisted in SS cells (CW: 45%, CCW: 31%) but was largely abolished in NS cells, which displayed nearly equal occupancy of the CW and CCW states (CW: 39%, CCW: 37%). Although pause-state occupancy remained lower than run-state occupancy in all conditions, it increased modestly under environmental O_2_, suggesting a greater contribution of pauses to overall motor-state residence under this oxygen condition.

The directionality of direct run-to-run transitions was modulated by both oxygen availability and magnetic polarity. Under low O_2_, SS cells preferentially switched from CW to CCW, whereas NS cells exhibited the opposite bias, favoring CCW to CW transitions. Under environmental O_2_, the directional bias was reversed in SS cells, while NS cells displayed nearly symmetric transition probabilities between the CW and CCW states. Together, these results show that although the overall switching architecture was conserved across conditions and dominated by direct CCW ↔ CW reversals, oxygen availability and magnetic polarity shaped both state occupancy and the preferred direction of motor-state transitions.

### 3.3 The free swimming dynamics follows similar framework as the tethered bacterial flagellar motor dynamics

Because tethering restricts the natural swimming behavior of MSR-1, we next analyzed free-swimming MSR-1 cells under reference-state conditions. To isolate the effects of oxygen availability while minimizing polarity-dependent variability, the analysis was restricted to SS cells in the absence of added chemical stimuli. Reconstructed trajectories were used to quantify swimming states, transition dynamics, state durations, and swimming speeds.

Free swimming of SS cells under reference-state conditions was characterized using a three-state semi-Markov chain comprising forward swimming, backward swimming, and pauses. For clarity, forward swimming is referred to here as the run state, whereas backward swimming represents the reverse-swimming or reversal state. Figure 7 shows that state residence was dominated by active swimming under both oxygen conditions, with pauses contributing only a negligible fraction of total occupancy. Under low O_2_, cells spent 71% of the time in the forward-run state and 29% in the reverse-swimming state, while pause occupancy was nearly negligible at 0.06%. Under environmental O_2_, this asymmetry became more pronounced, with cells spending 84% of the time in the forward-run state and 16% in the reverse-swimming state, while pause occupancy remained very low at 0.09%. Because state occupancy reflects both the frequency of state entry and the duration of state residence, the reduced occupancy of the reverse-swimming state does not necessarily imply fewer reversal events. Instead, when considered alongside the nearly balanced transition probabilities between forward and reverse swimming, it indicates that reversals were shorter and/or forward runs were longer under environmental O_2_. Despite these differences in residence time, the transition network remained dominated by direct run–reversal exchanges, with pause-mediated transitions representing a secondary pathway. Although pauses occurred more frequently under environmental O_2_, they remained transient intermediate states from which cells rapidly resumed active swimming. Together, these results indicate that reference-state free swimming in SS MSR-1 is organized primarily as a run-reversal process, with pauses acting as rare, condition-dependent intermediate states rather than as a major residence state. The agreement between tethered and free-swimming measurements further indicates that tethered-cell assays provide a sensitive motor-level readout of motility adaptation to environmental cues.

**Figure 7.**
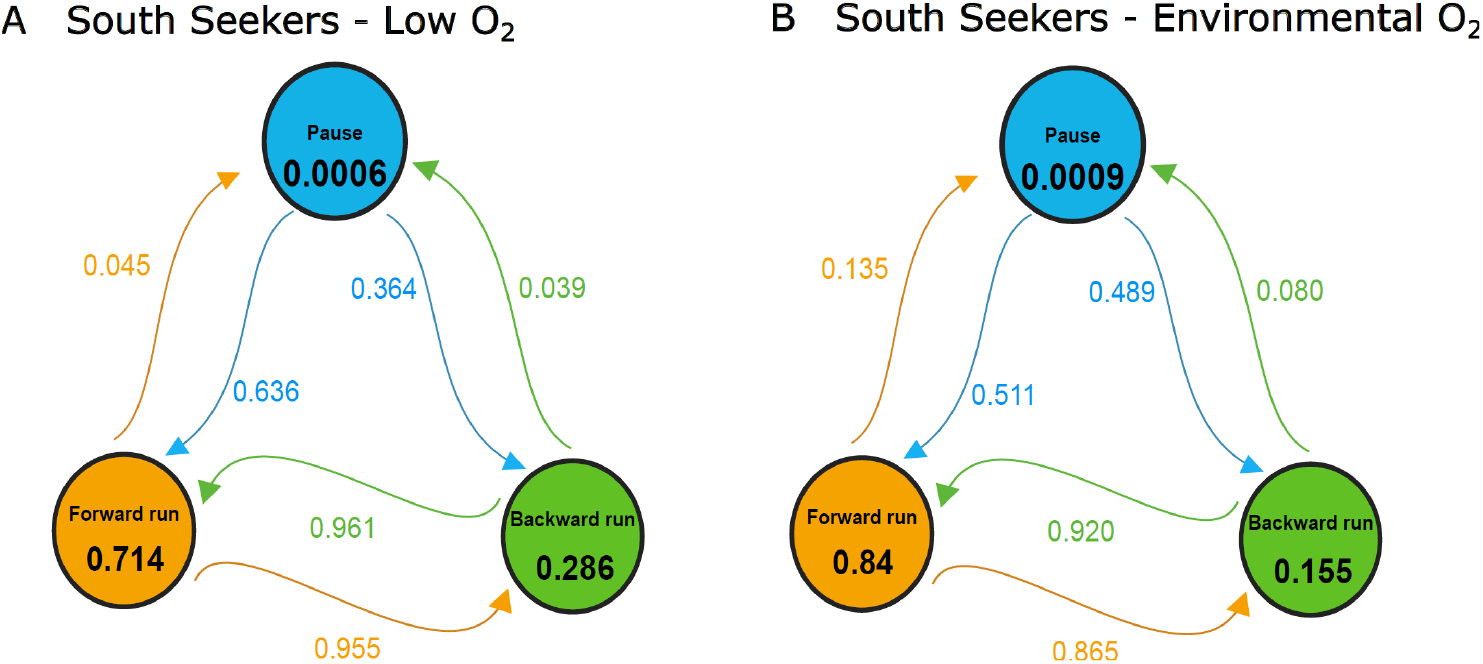
Semi-Markov chain model for free-swimming MSR-1 cells. Semi-Markov chains showing the transition probabilities and state occupancies of forward runs, backward runs, and pauses in south-seeking cells under low O_2_ (A) and environmental O_2_ (B) conditions. Node values indicate relative state occupancy, and arrows indicate transition probabilities.

## 4 Discussion

Bacterial motility for MTB cells like MSR-1 integrates multiple sensory inputs, therefore to understand the BFM chemotactic response it is necessary to first characterize the baseline motor dynamics under reference-state conditions to distinguish between aerotaxis, magnetotaxis and chemotaxis. The quantitative model of the reference state provides a foundation for elucidating chemotactic responses at the molecular level through BFM dynamics and for understanding how these dynamics are integrated with the broader chemotaxis system and other forms of taxis.

### 4.1 MSR-1 motility exhibits non-Markovian temporal organization

State residence time revealed a critical feature of the motor dynamics: pause durations, time between switches, and free-swimming runs durations were better described by lognormal-like distributions than by exponential distributions. This finding is important because exponential dwell-time distributions arise from memoryless Poisson processes, in which the probability of leaving a state is constant and independent of the time already spent in that state [15]. The preferential fit of lognormal-like distributions therefore suggests that the probability of leaving a swimming state depends on past events and are influenced by underlying biological or mechanical processes rather than occuring at constant rate [30]. Similar non-Poissonian switching has been reported for polar-flagellated bacteria such as *Vibrio alginolyticus*, where non-exponential forward and backward interval distributions were interpreted as evidence of refractory or history-dependent motor switching [46, 45]. In MSR-1, the same principle appears to apply at both the tethered-motor and free-swimming levels: the observed states are not simply Markovian states with constant transition rates, but coarse-grained outputs of the canonical chemotaxis pathway, and the dual motor synchronization. This required the use of a semi-Markov framework, where the embedded transition matrix captures the order of state transitions, while the dwell-time distributions capture the temporal persistence of each state.

### 4.2 A conserved reversal-dominated architecture organizes MSR-1 motility

Analysis of tethered cells under reference-state conditions revealed that direct transitions between CCW and CW rotation constitute the primary motor-switching pathway, irrespective of oxygen concentration or magnetic polarity. Although pause-mediated transitions represented only a minority of switching events, their consistent occurrence across all conditions indicates that pausing is an intrinsic and reproducible component of MSR-1 motor dynamics. This finding agrees with previous studies identifying pauses as a genuine third motor state rather than an experimental artifact or a transient cessation of rotation [49, 21]. Importantly, while pauses do not dominate the overall switching architecture, their reproducible occurrence means they cannot be neglected when describing motor-state dynamics. Their inclusion as a third state provides a more complete representation of the MSR-1 flagellar motor than the conventional two-state model.

Across all conditions, direct CCW ↔ CW reversals remained the dominant switching mechanism, whereas pause-mediated transitions constituted a reproducible secondary pathway. In MSR-1, these pauses occurred in two distinct transition contexts: the BFM either resumed rotation in the same direction as before the pause or in the opposite direction. Interestingly, direction-retaining pauses were consistently more frequent than direction-switching pauses. However, the similar duration of these two classes suggest that they may arise from a common or closely related paused motor state, while differing in how that state is resolved. This interpretation is consistent with previous studies linking BFM pausing to components of the chemotaxis signaling and motor-switching machinery [21, 10, 9]. Direction-retaining pauses have been proposed to represent incomplete or futile switching attempts in which the motor enters a switching-related intermediate state but ultimately returns to its original rotational direction. These transient pauses that do not culminate in reversal have also been described in other bacterial systems supporting its classification as a type of pause state [26]. In contrast, direction-switching pauses can be associated with a direction change in the BFM that spreads to the flagellar configuration promoting a change in position to accomodate the new direction, this has been observed by Murat et al., showing that in the bipolarly flagellated bacteria *Magnetospirillum magneticum* (AMB-1), reversal is unlikely to result solely from a change in rotor rotational state. Rather, reversal must also be coordinated with changes in the activity and configuration of the opposite polar flagella. In AMB-1, the flagella can switch between different configurations by wrapping around or unwrapping from the cell body, thereby altering cell motility [27]. A similar pattern was reported by Pfeiffer et al. where MSR-1 undergoes into transient pauses during transitions between wrapped and unwrapped flagellar configurations, and that these transitions are associated with motor-direction switching since the cell body direction changes from the previous trajectory [28]. Together, these observations suggest that pauses associated with reversals represent direction-switching states, rather than simple interruptions of motility, coupling motor reversal with flagellar reconfiguration, including flagellar wrapping or unwrapping, and coordination between the two polar motors.

Extending the analysis from tethered motors to freely swimming cells demonstrated that the reversal-dominated architecture was preserved at the whole-cell level. This conservation suggests that the reversal-dominated switching pattern observed in tethered assays reflects an intrinsic feature of MSR-1 motility rather than an organization imposed by mechanical tethering. The persistence of non-Poissonian dwell-time distributions further indicates that the temporal organization of motility is maintained across experimental configurations. Importantly, no complete pauses were observed during free-swimming experiments, supporting the idea that pauses detected at the level of individual motors may correspond to flagellar wrapping or unwrapping events rather than complete cessation of cell motility. Such events could facilitate reorientation of the cell body and may account for the transient decrease in swimming speed frequently observed before a change in swimming direction. This interpretation suggests that the reduction in swimming speed may reflect the temporary cessation or slowing of one motor, allowing flagellar reconfiguration between wrapped and unwrapped states.

### 4.3 Magnetic polarity biases transition pathways, whereas oxygen modulates state persistence

Analysis of pause dynamics showed that oxygen availability modulated pausing primarily through changes in the frequency and duration of pause events. Unlike classical aerotaxis, which is driven by spatial oxygen gradients [31, 12], our results indicate that BFM dynamics remain sensitive to oxygen availability even under nominally uniform oxygen conditions. This interpretation is consistent with observations in *Magnetovirga frankeli* (SS-5), where sustained exposure to low oxygen reduced swimming speed [13]. Moreover, the polarity dependence of these effects suggests that oxygen-dependent regulation is conditioned by the magnetic and swimming polarity of the cell, potentially through asymmetric control or coordination of the two polar flagellar motors. Rather than demonstrating a separate magnetotactic signaling pathway, these findings support functional coupling between oxygen sensing and the polarity-dependent organization of the MSR-1 motility system.

The semi-Markov analysis revealed a hierarchical organization of MSR-1 motor dynamics. At its core, the system is dominated by direct CCW↔CW reversals that are largely invariant across oxygen conditions and magnetic polarity. Superimposed on this conserved switching architecture is a second regulatory layer formed by pause states, whose entry pathway depends primarily on cell polarity, whereas their residence time is modulated by oxygen availability.

In freely swimming cells, oxygen had only a limited influence on transition probabilities, leaving the reversal-dominated architecture largely unchanged, it more substantially altered the relative occupancy of the motility states. Pause-like states detected during free swimming should not be interpreted as complete motor arrests, but rather as transient low-speed states within the reversal sequence. Together, these findings indicate that oxygen primarily modulates the temporal organization and relative occupancy of motility states while preserving the underlying reversal architecture.

These results support a hierarchical model of MSR-1 motility in which a conserved reversal-dominated switching architecture is modulated by two complementary regulatory mechanisms: magnetic polarity biases the pathways through which motor-state transitions occur, whereas oxygen availability primarily governs the temporal organization of those states by modulating their residence times.

## 5 Conclusions

This study establishes a quantitative reference-state model for MSR-1 flagellar-motor and wholecell motility dynamics. The results support a semi-Markov description in which a conserved reversal-dominated architecture forms the core of the motility profile, while pause states constitute reproducible intermediate pathways that contribute to motor switching and whole-cell reorientation. The preservation of this organization across tethered and freely swimming cells indicates that it represents an intrinsic feature of MSR-1 motility rather than an artifact of the experimental configuration.

Oxygen availability and magnetic polarity modulate distinct but complementary dimensions of this architecture. Magnetic polarity biases the pathways through which pause-associated transitions are accessed, whereas oxygen primarily influences the temporal persistence and relative occupancy of motility states. MSR-1 motility should therefore be understood not as a fixed un-stimulated state, but as a condition-dependent dynamical regime in which a stable reversal core is differentially modulated by environmental and polarity-dependent inputs. This framework provides a defined baseline for identifying how future chemical stimuli perturb motor-state transitions, residence times, and whole-cell swimming behavior.

## References

[1] Dennis A. Bazylinski and Timothy J. Williams. Ecophysiology of Magnetotactic Bacteria. In Dirk Schüler, editor, Magnetoreception and Magnetosomes in Bacteria, pages 37–75. Springer, Berlin, Heidelberg, 2007.

[2] Howard C. Berg and Douglas A. Brown. Chemotaxis in Escherichia coli analysed by Three-dimensional Tracking. Nature, 239(5374):500–504, October 1972.

[3] Richard Blakemore. Magnetotactic bacteria. Science, 190(4212):377–379, 1975.

[4] Steven M. Block, Jeffrey E. Segall, and Howard C. Berg. Impulse responses in bacterial chemotaxis. Cell, 31(1):215–226, November 1982.

[5] Thierry Darnige, Daniel Midtvedt, Renaud Baillou, Benjamin Perez Estay, Changsong Wu, Alex Le Guen, Giovanni Volpe, and Eric Clement. Deep learning-enhanced Lagrangian 3D Tracking of motile microorganisms, March 2026.

[6] Edward F. DeLong, Richard B. Frankel, and Dennis A. Bazylinski. Multiple evolutionary origins of magnetotaxis in bacteria. Science, 259(5096):803–806, 1993.

[7] Mariia Dvoriashyna and Eric Lauga. Hydrodynamics and direction change of tumbling bacteria. PLOS ONE, 16(7):e0254551, 2021.

[8] Michael Eisenbach. Control of bacterial chemotaxis. Molecular Microbiology, 20(5):903–910, 1996.

[9] Michael Eisenbach, Amnon Wolf, Martin Welch, S. Roy Caplan, I. R. Lapidus, Robert M. Macnab, Hamutal Aloni, and Ora Asher. Pausing, switching and speed fluctuation of the bacterial flagellar motor and their relation to motility and chemotaxis. Journal of Molecular Biology, 211(3):551–563, 1990.

[10] Michael Eisenbachlg, Amnon WolfI, Martin WelchI, S Roy Caplanl, I Richard, Robert M Macnab, Hamutal Alonil, and Ora Asher’. Pausing, Switching and Speed Fluctuation of the Bacterial Flagellar Motor and their Relation to Motility and Chemotaxisj-.

[11] Damien Faivre and Dirk Schüler. Magnetotactic bacteria and magnetosomes. Chemical Reviews, 108(11):4875–4898, 2008.

[12] R. B. Frankel, D. A. Bazylinski, M. S. Johnson, and B. L. Taylor. Magneto-aerotaxis in marine coccoid bacteria. Biophysical Journal, 73(2):994–1000, August 1997.

[13] Emilie Gachon, Sascha Lambert, Sandrine Grosse, Elsa Turrini, Emma Ropion, Stefan Klumpp, Christopher T. Lefèvre, Mila Sirinelli, and Damien Faivre. A magnetotactic bacterium capable of magnetic sensing. iScience, 28(9):113377, 2025.

[14] Y A Gorby, T J Beveridge, and R P Blakemore. Characterization of the bacterial magnetosome membrane. Journal of Bacteriology, 170(2):834–841, February 1988.

[15] Moshe Haviv. The exponential distribution and the poisson process. In Queues: A Course in Queueing Theory, pages 1–19. Springer, New York, NY, 2013.

[16] Basarab G. Hosu, Vedavalli S. J. Nathan, and Howard C. Berg. Internal and external components of the bacterial flagellar motor rotate as a unit. Proceedings of the National Academy of Sciences, 113(17):4783–4787, April 2016.

[17] Steven Johnson, Justin C. Deme, Emily J. Furlong, Joseph J. E. Caesar, Fabienne F. V. Chevance, Kelly T. Hughes, and Susan M. Lea. Structural basis of directional switching by the bacterial flagellum. Nature Microbiology, 9(5):1282–1292, 2024.

[18] Steven Johnson, Emily J. Furlong, Justin C. Deme, Ashley L. Nord, Joseph J. E. Caesar, Fabienne F. V. Chevance, Richard M. Berry, Kelly T. Hughes, and Susan M. Lea. Molecular structure of the intact bacterial flagellar basal body. Nature Microbiology, 6(6):712–721, 2021.

[19] R. Killick, P. Fearnhead, and I. A. Eckley. Optimal Detection of Changepoints With a Linear Computational Cost. Journal of the American Statistical Association, 107(500):1590–1598, December 2012.

[20] Mila Kojadinovic, Antoine Sirinelli, George H. Wadhams, and Judith P. Armitage. New motion analysis system for characterization of the chemosensory response kinetics of Rhodobacter sphaeroides under different growth conditions. Applied and Environmental Microbiology, 77(12):4082–4088, 2011.

[21] I. R. Lapidus, M. Welch, and M. Eisenbach. Pausing of flagellar rotation is a component of bacterial motility and chemotaxis. Journal of Bacteriology, 170(8):3627–3632, 1988.

[22] Christopher T. Lefèvre and Dennis A. Bazylinski. Ecology, diversity, and evolution of magnetotactic bacteria. Microbiology and Molecular Biology Reviews, 77(3):497–526, 2013.

[23] Christopher T. Lefèvre, Mathieu Bennet, Livnat Landau, Peter Vach, David Pignol, Dennis A. Bazylinski, Richard B. Frankel, Stefan Klumpp, and Damien Faivre. Diversity of Magneto-Aerotactic Behaviors and Oxygen Sensing Mechanisms in Cultured Magnetotactic Bacteria. Biophysical Journal, 107(2):527–538, July 2014.

[24] Yukio Magariyama, Makoto Ichiba, Kousou Nakata, Kensaku Baba, Toshio Ohtani, Seishi Kudo, and Tomonobu Goto. Difference in bacterial motion between forward and backward swimming caused by the wall effect. Biophysical Journal, 88(5):3648–3658, 2005.

[25] Xuegang Mao, Ramon Egli, Nikolai Petersen, and Xiuming Liu. Combined response of polar magnetotaxis to oxygen and pH: Insights from hanging drop assays and microcosm experiments. Scientific Reports, 14(1):27331, 2024.

[26] Tanmoy Mukherjee, Mustafa Elmas, Lam Vo, Vasilios Alexiades, Tian Hong, and Gladys Alexandre. Multiple CheY Homologs Control Swimming Reversals and Transient Pauses in Azospirillum brasilense. Biophysical Journal, 116(8):1527–1537, April 2019.

[27] Dorothée Murat, Marion Hérisse, Leon Espinosa, Alicia Bossa, François Alberto, and Long-Fei Wu. Opposite and coordinated rotation of amphitrichous flagella governs oriented swimming and reversals in a magnetotactic spirillum. Journal of Bacteriology, 197(20):3275–3282, 2015.

[28] Lena Oertwig and Daniel Pfeiffer. Insights into unique flagellar motor dynamics in an amphitrichously flagellated magnetotactic spirillum. Conference presentation, VAAM 2026, session “Prokaryotic Cell Biology”, 2026. Conference abstract; no journal article or DOI was identified.

[29] Teuta Pilizota, Mostyn T. Brown, Mark C. Leake, Richard W. Branch, Richard M. Berry, and Judith P. Armitage. A molecular brake, not a clutch, stops the Rhodobacter sphaeroides flagellar motor. Proceedings of the National Academy of Sciences of the United States of America, 106(28):11582–11587, 2009.

[30] Jennifer Pohle, Timo Adam, and Larissa T. Beumer. Flexible estimation of the state dwell-time distribution in hidden semi-markov models. Computational Statistics & Data Analysis, 172:107479, 2022.

[31] Felix Popp, Judith P. Armitage, and Dirk Schüler. Polarity of bacterial magnetotaxis is controlled by aerotaxis through a common sensory pathway. Nature Communications, 5:5398, 2014.

[32] M. Reufer, R. Besseling, J. Schwarz-Linek, V. A. Martinez, A. N. Morozov, J. Arlt, D. Trubitsyn, F. B. Ward, and W. C. K. Poon. Switching of swimming modes in Magnetospirillium gryphiswaldense. Biophysical Journal, 106(1):37–46, 2014.

[33] Sheri L. Simmons and Katrina J. Edwards. Geobiology of Magnetotactic Bacteria. In Dirk Schüler, editor, Magnetoreception and Magnetosomes in Bacteria, pages 77–102. Springer, Berlin, Heidelberg, 2007.

[34] Stefan Spring, Rudolf Amann, Wolfgang Ludwig, Karl-Heinz Schleifer, Hans Van Gemerden, and Nikolai Petersen. Dominating Role of an Unusual Magnetotactic Bacterium in the Microaerobic Zone of a Freshwater Sediment. Applied and Environmental Microbiology, 59(8):2397–2403, August 1993.

[35] Stefan Spring and Dennis A. Bazylinski. Magnetotactic bacteria. In The Prokaryotes, pages 842–862. Springer, New York, NY, 2006.

[36] John F. Stolz. Magnetosomes. Microbiology, 139(8):1663–1670, 1993.

[37] Jiaxing Tan, Ling Zhang, Xingtong Zhou, Siyu Han, Yan Zhou, and Yongqun Zhu. Structural basis of the bacterial flagellar motor rotational switching. Cell Research, 34(11):788–801, 2024.

[38] Hiroto Tanaka, Yasuaki Kazuta, Yasushi Naruse, Yukihiro Tominari, Hiroaki Umehara, Yoshiyuki Sowa, Takashi Sagawa, Kazuhiro Oiwa, Masato Okada, Ikuro Kawagishi, and Hiroaki Kojima. Bayesian-based decipherment of in-depth information in bacterial chemical sensing beyond pleasant/unpleasant responses. Scientific Reports, 12(1):2965, February 2022.

[39] Linda Turner, William S. Ryu, and Howard C. Berg. Real-Time Imaging of Fluorescent Flagellar Filaments. Journal of Bacteriology, 182(10):2793–2801, May 2000.

[40] René Uebe and Dirk Schüler. Magnetosome biogenesis in magnetotactic bacteria. Nature Reviews Microbiology, 14(10):621–637, October 2016.

[41] George H. Wadhams and Judith P. Armitage. Making sense of it all: Bacterial chemotaxis. Nature Reviews Molecular Cell Biology, 5(12):1024–1037, 2004.

[42] Carina Weigel and Daniel Pfeiffer. Directional matching of swimming polarity provides a competitive advantage during bacterial magneto-aerotaxis. BMC Microbiology, 26(1):449, May 2026.

[43] Changsong Wu. Aerotatic Response and Magnetic Control of a Magnetotactic Bacterium (MSR-1). PhD thesis, Sorbonne Université, Paris, France, September 2025.

[44] Bohan Wu-Zhang, Peixin Zhang, Renaud Baillou, Anke Lindner, Eric Clément, Gerhard Gompper, and Dmitry A. Fedosov. Run-and-tumble dynamics of Escherichia coli is governed by its mechanical properties. Journal of the Royal Society Interface, 22(227):20250035, 2025.

[45] Li Xie, Tuba Altindal, and Xiao-Lun Wu. An element of determinism in a stochastic flagellar motor switch. PLOS ONE, 10(11):e0141654, 2015.

[46] Yang Yang, Jing He, Tuba Altindal, Li Xie, and Xiao-Lun Wu. A non-poissonian flagellar motor switch increases bacterial chemotactic potential. Biophysical Journal, 109(5):1058–1069, 2015.

[47] Wei-Jia Zhang and Long-Fei Wu. Flagella and swimming behavior of marine magnetotactic bacteria. Biomolecules, 10(3):460, 2020.

[48] Xiaotian Zhou and Anna Roujeinikova. The structure, composition, and role of periplasmic stator scaffolds in polar bacterial flagellar motors. Frontiers in Microbiology, 12:639490, 2021.

[49] Xiang-Yu Zhuang and Chien-Jung Lo. Decoding bacterial motility: From swimming states to patterns and chemotactic strategies. Biomolecules, 15(2):170, 2025.

