## Supplemental Data for "From flagellar motor behavior to bacterial swimming: defining a reference state for motility dynamics in *Magnetospirillum gryphiswaldense*"

September 9, 2026

### Supplemental Information

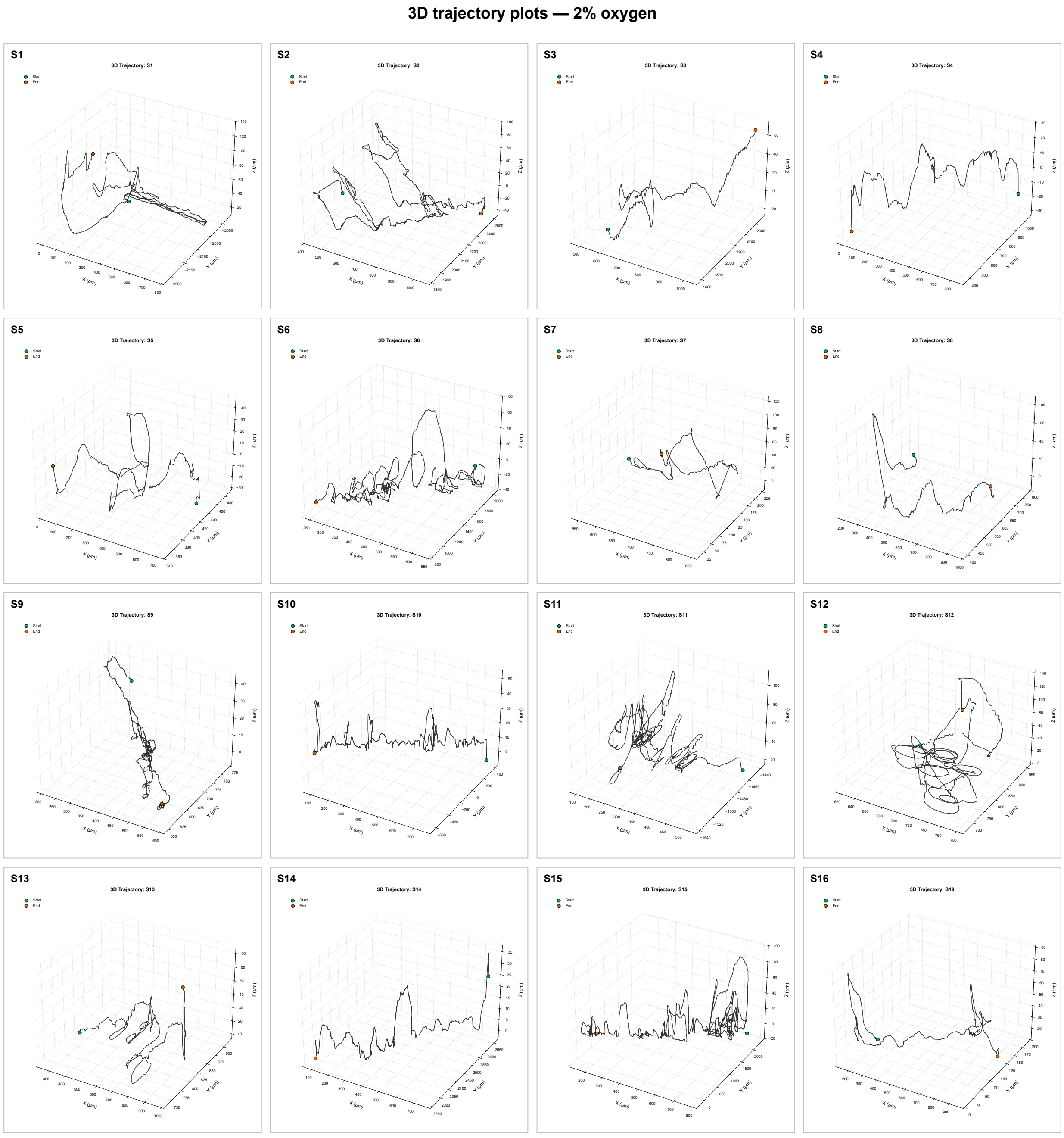


Figure 1: Three-dimensional free-swimming trajectories of south-seeking MSR-1 cells under Low $O_2$ conditions. Each panel shows one reconstructed cell trajectory, with the black line representing the swimming path in 3D space. The $x$, $y$, and $z$ axes are reported in $\mu$m. Green and orange markers indicate the start and end positions of each trajectory, respectively.

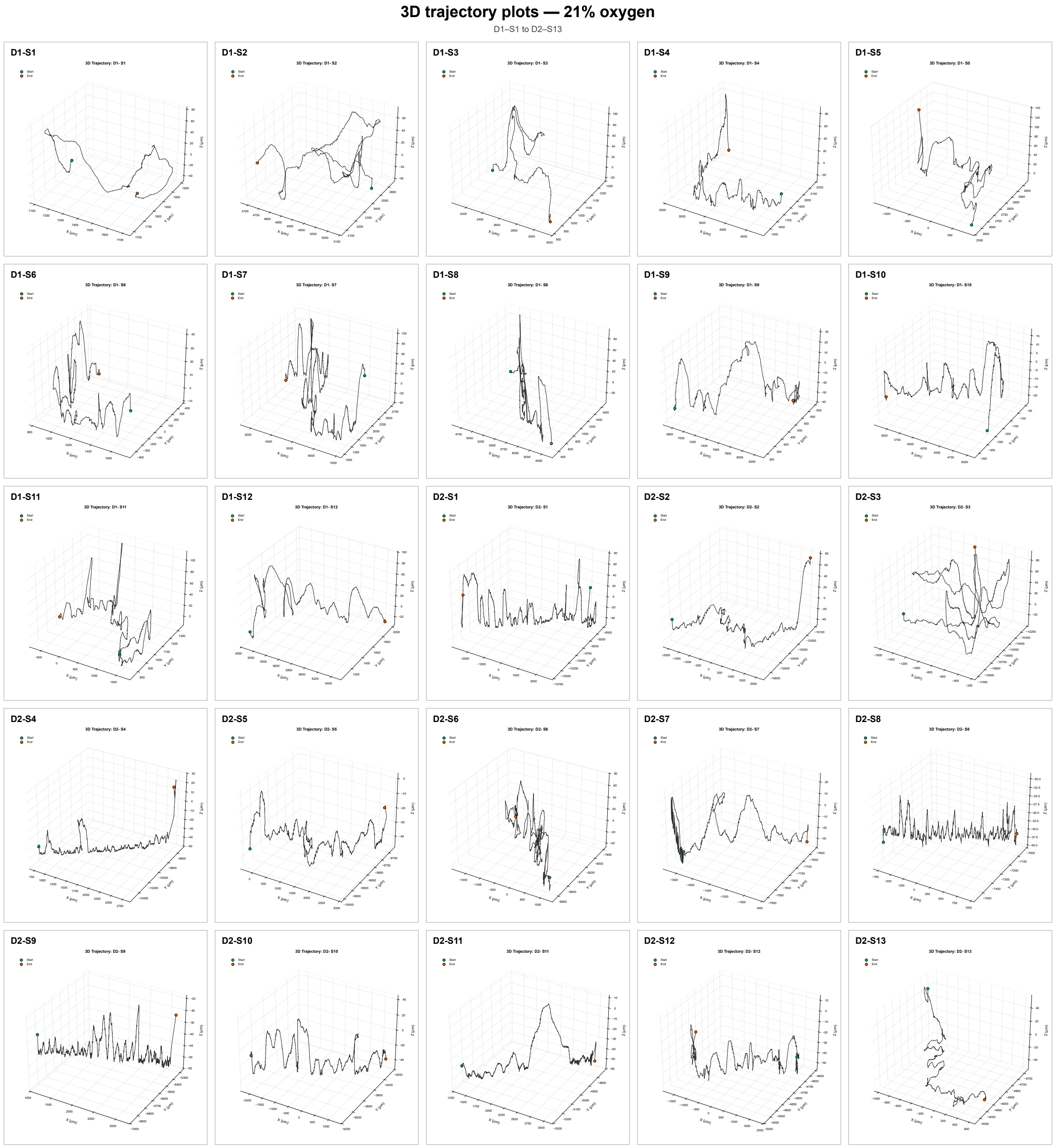


Figure 2: Three-dimensional free-swimming trajectories of south-seeking MSR-1 cells under environmental $O_2$ conditions. Each panel shows one reconstructed cell trajectory, with the black line representing the swimming path in 3D space. The $x$, $y$, and $z$ axes are reported in $\mu$m. Green and orange markers indicate the start and end positions of each trajectory, respectively

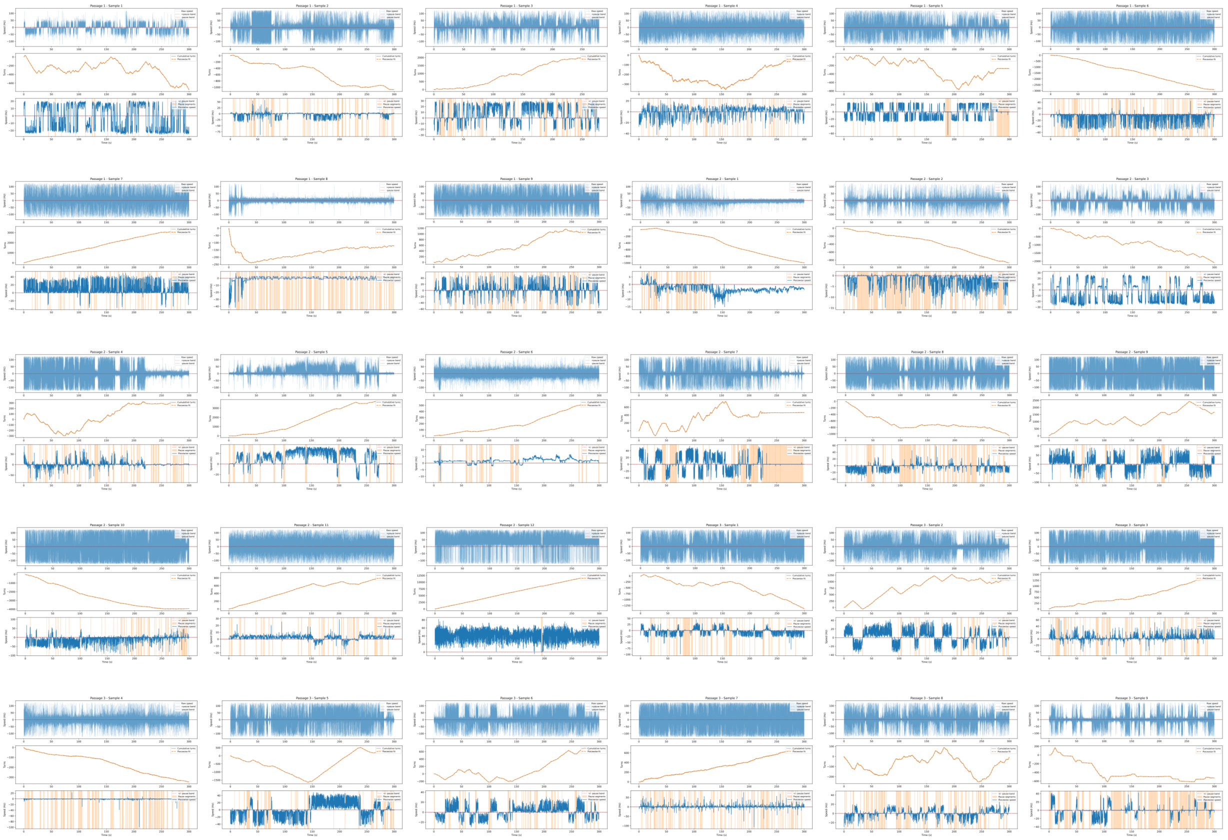

Figure 3: Plots for tethered-cell rotational trajectories for South Seekers under the low $O_2$ condition. Each panel corresponds to one analyzed cell trajectory, ordered by passage number and sample number. For each sample, the upper subplot shows the raw instantaneous rotational speed over time, with the pause-speed band indicated around zero. The middle subplot shows the cumulative number of turns as a function of time together with the corresponding piecewise fit used to describe changes in rotational behavior. The lower subplot shows the piecewise-estimated rotational speed, with shaded vertical regions marking intervals classified as pause-like events.

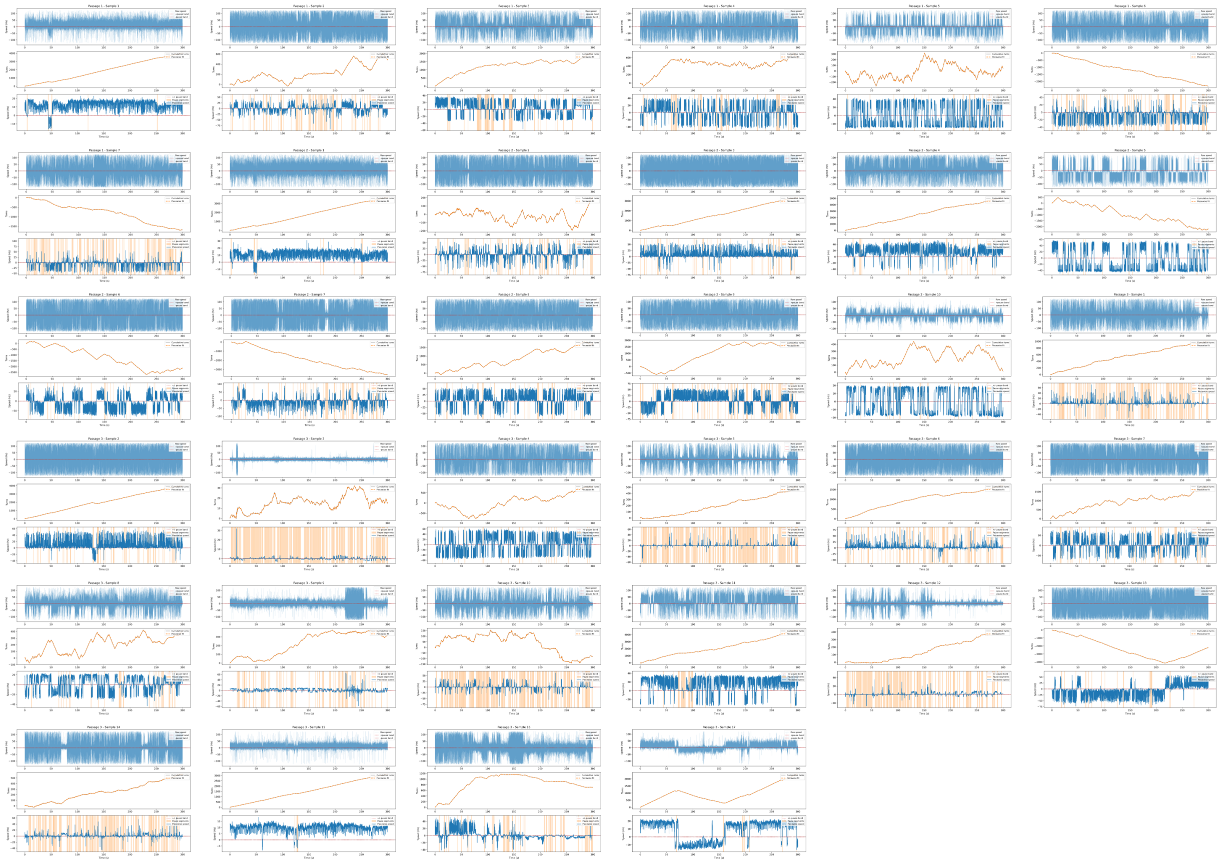

Figure 4: Plots for tethered-cell rotational trajectoriesfor North Seekers under the low $O_2$ condition. Each panel corresponds to one analyzed cell trajectory, ordered by passage number and sample number. For each sample, the upper subplot shows the raw instantaneous rotational speed over time, with the pause-speed band indicated around zero. The middle subplot shows the cumulative number of turns as a function of time together with the corresponding piecewise fit used to describe changes in rotational behavior. The lower subplot shows the piecewise-estimated rotational speed, with shaded vertical regions marking intervals classified as pause-like events.

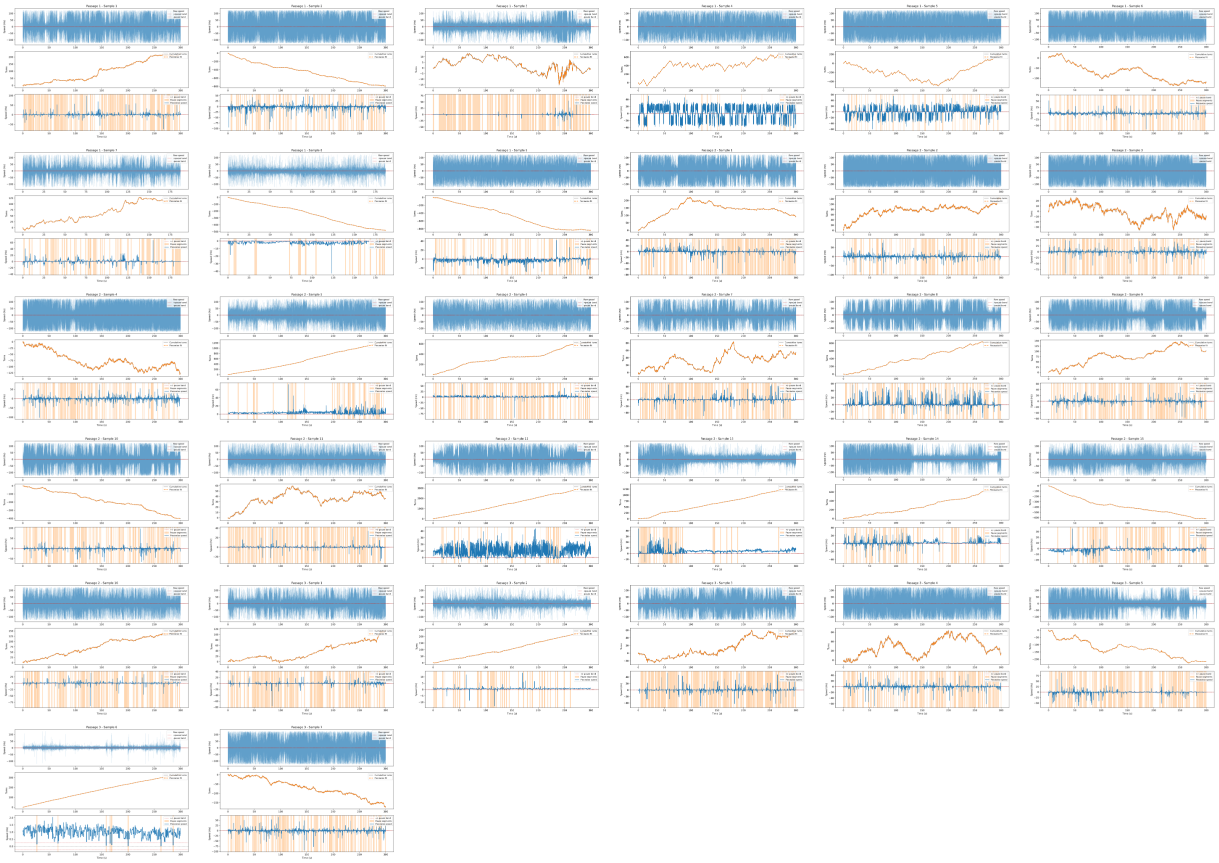

Figure 5: Plots for tethered-cell rotational trajectories for South Seekers under the environmental $O_2$ condition. Each panel corresponds to one analyzed cell trajectory, ordered by passage number and sample number. For each sample, the upper subplot shows the raw instantaneous rotational speed over time, with the pause-speed band indicated around zero. The middle subplot shows the cumulative number of turns as a function of time together with the corresponding piecewise fit used to describe changes in rotational behavior. The lower subplot shows the piecewise-estimated rotational speed, with shaded vertical regions marking intervals classified as pause-like events.

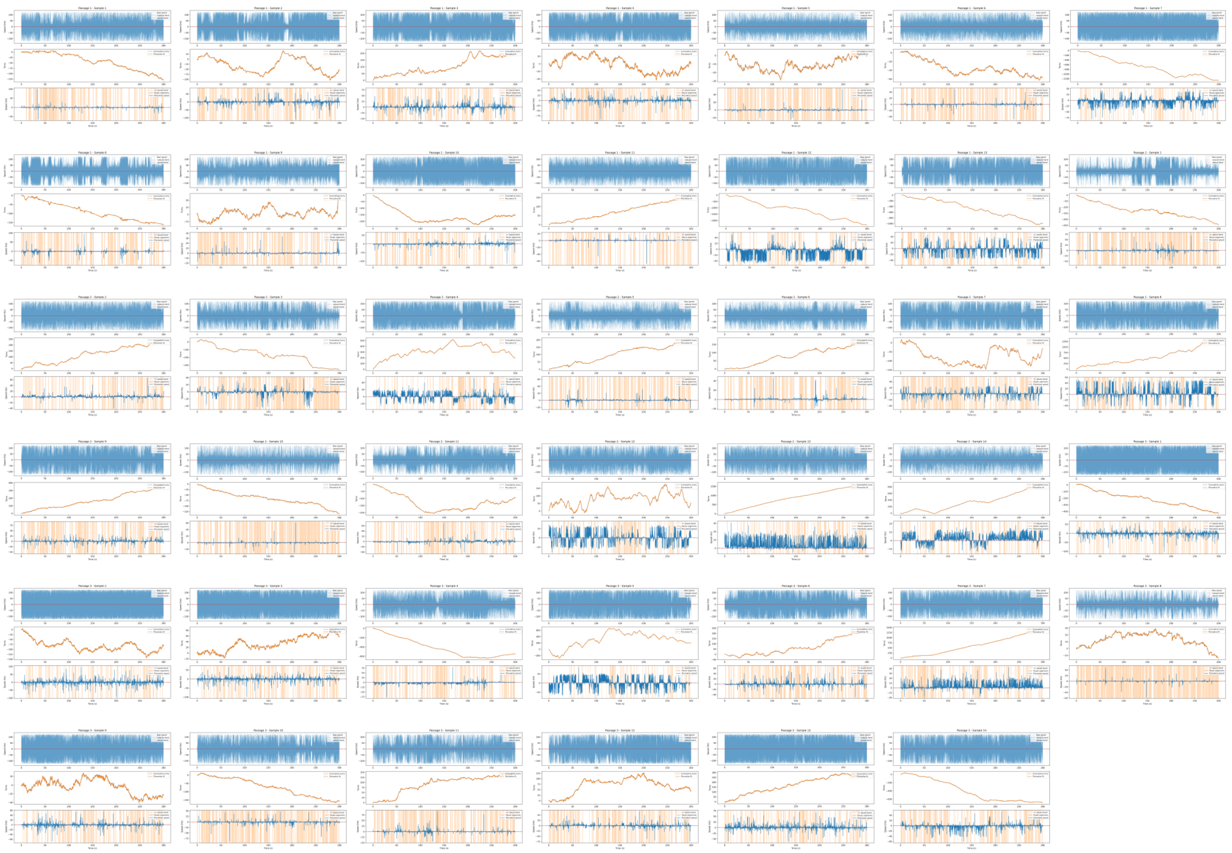

Figure 6: Plots for tethered-cell rotational trajectories for North Seekers under the environmental $O_2$ condition. Each panel corresponds to one analyzed cell trajectory, ordered by passage number and sample number. For each sample, the upper subplot shows the raw instantaneous rotational speed over time, with the pause-speed band indicated around zero. The middle subplot shows the cumulative number of turns as a function of time together with the corresponding piecewise fit used to describe changes in rotational behavior. The lower subplot shows the piecewise-estimated rotational speed, with shaded vertical regions marking intervals classified as pause-like events.

Table S1: Goodness-of-fit statistics for candidate distributions fitted to pause durations and inter-switch times under low $O_2$ and environmental $O_2$ conditions. Values correspond to Kolmogorov–Smirnov (KS) statistics; lower values indicate better agreement between the empirical data and the fitted distribution. The best-fitting distribution within each condition is highlighted in bold.

| Variable | Distribution | Low $O_2$ | | Environmental $O_2$ | |
|---|---|---|---|---|---|
| | | NS | SS | NS | SS |
| Pause duration | Normal | 0.23972 | 0.27740 | 0.17090 | 0.16886 |
| | Lognormal | **0.05109** | **0.04457** | **0.02856** | **0.02857** |
| | Weibull | 0.10245 | 0.11818 | 0.06967 | 0.07042 |
| | Exponential | 0.11667 | 0.11858 | 0.13435 | 0.13624 |
| | Gamma | 0.10848 | 0.12318 | 0.06915 | 0.06350 |
| Time between Switches | Normal | 0.33745 | 0.37858 | 0.25273 | 0.28374 |
| | Lognormal | **0.04579** | **0.06364** | **0.03300** | **0.02809** |
| | Weibull | 0.10190 | 0.11728 | 0.09885 | 0.10472 |
| | Exponential | 0.17402 | 0.20949 | 0.09865 | 0.09499 |
| | Gamma | 0.11105 | 0.14437 | 0.10045 | 0.10274 |

Table S2: Lognormal parameter estimates for pause durations and inter-switch times under environmental $O_2$ and low $O_2$ conditions. Estimates are reported with their 95% confidence intervals in brackets. SD refers to the standard deviation of the parameter estimate.

| Variable | Seekers | $O_2$ condition | Parameter | Estimate [95% CI] | SD |
|---|---|---|---|---|---|
| Pause duration | South | Environmental $O_2$ | Location $\mu$ | -0.617 [-0.646, -0.587] | 0.015 |
| | | | Scale $\sigma$ | 0.816 [0.795, 0.837] | 0.011 |
| | North | Environmental $O_2$ | Location $\mu$ | -0.663 [-0.688, -0.638] | 0.013 |
| | | | Scale $\sigma$ | 0.822 [0.805, 0.840] | 0.009 |
| | South | Low $O_2$ | Location $\mu$ | -0.633 [-0.677, -0.589] | 0.022 |
| | | | Scale $\sigma$ | 0.888 [0.857, 0.920] | 0.016 |
| | North | Low $O_2$ | Location $\mu$ | -0.793 [-0.834, -0.752] | 0.021 |
| | | | Scale $\sigma$ | 0.884 [0.855, 0.914] | 0.015 |
| Time between switches | South | Environmental $O_2$ | Location $\mu$ | -1.033 [-1.056, -1.011] | 0.012 |
| | | | Scale $\sigma$ | 0.943 [0.927, 0.959] | 0.008 |
| | North | Environmental $O_2$ | Location $\mu$ | -1.042 [-1.061, -1.024] | 0.009 |
| | | | Scale $\sigma$ | 0.945 [0.932, 0.958] | 0.007 |
| | South | Low $O_2$ | Location $\mu$ | -0.673 [-0.712, -0.634] | 0.020 |
| | | | Scale $\sigma$ | 1.205 [1.177, 1.233] | 0.014 |
| | North | Low $O_2$ | Location $\mu$ | -0.539 [-0.575, -0.503] | 0.018 |
| | | | Scale $\sigma$ | 1.236 [1.210, 1.262] | 0.013 |

Table S3: Goodness-of-fit statistics for candidate dwell-time distributions fitted to the 3D motility states under low $O_2$ and environmental $O_2$ conditions. For each state and condition, the number of events (*N*), Bayesian information criterion (BIC), and Kolmogorov–Smirnov statistic (KS) are reported. Lower BIC and KS values indicate better fit. The lowest BIC and lowest KS value within each state and condition are highlighted in bold.

| $O_2$ condition | State | Distribution | *N* | BIC | KS statistic |
|---|---|---|---|---|---|
| Low $O_2$ | Forward | Exponential | 274 | 1261.306 | 0.193 |
| | | Gamma | 274 | 1201.833 | 0.102 |
| | | Lognormal | 274 | **1167.471** | **0.049** |
| | | Weibull | 274 | 1184.871 | 0.065 |
| | Pause | Exponential | 22 | -94.790 | 0.610 |
| | | Gamma | 22 | -168.930 | 0.500 |
| | | Lognormal | 22 | **-170.292** | **0.471** |
| | | Weibull | 22 | -159.556 | 0.491 |
| | Backward | Exponential | 268 | 754.687 | 0.129 |
| | | Gamma | 268 | 751.471 | 0.082 |
| | | Lognormal | 268 | **716.953** | **0.070** |
| | | Weibull | 268 | 742.788 | 0.075 |
| Environmental $O_2$ | Forward | Exponential | 223 | 1442.607 | 0.306 |
| | | Gamma | 223 | 1310.802 | 0.115 |
| | | Lognormal | 223 | **1262.983** | **0.056** |
| | | Weibull | 223 | 1283.598 | 0.074 |
| | Pause | Exponential | 45 | -175.311 | 0.526 |
| | | Gamma | 45 | -219.784 | 0.366 |
| | | Lognormal | 45 | **-234.868** | **0.352** |
| | | Weibull | 45 | -199.652 | 0.371 |
| | Backward | Exponential | 214 | 675.701 | 0.146 |
| | | Gamma | 214 | 656.807 | 0.099 |
| | | Lognormal | 214 | **621.066** | **0.052** |
| | | Weibull | 214 | 642.995 | 0.067 |

Table S4: Summary of pause occurrence and pause-duration statistics in free-swimming SS MSR-1 cells under low $O_2$ and environmental $O_2$ conditions. The sample-level pause fraction was calculated for each track as pause events divided by total events and is reported as the median with interquartile range, Q1–Q3. Pause-duration statistics were obtained from the semi-Markov statedwell-time.

**A. Pause occurrence**

| $O_2$ condition | Tracks | Windows | Total events | Pause events | Pooled pause fraction | Sample pause fraction [Q1–Q3] |
|---|---|---|---|---|---|---|
| Low $O_2$ | 16 | 65 | 565 | 22 | 3.89% | 2.10% [0.00–7.00] |
| Environmental $O_2$ | 25 | 110 | 485 | 45 | 9.28% | 6.25% [0.00–13.16] |

**B. Pause-duration structure**

| $O_2$ condition | Pause occupancy | Mean duration (s) | Median duration (s) | IQR duration (s) | SD duration (s) | CV duration |
|---|---|---|---|---|---|---|
| Low $O_2$ | 0.063% | 0.040 | 0.0375 | 0.0000 | 0.0049 | 0.124 |
| Environmental $O_2$ | 0.093% | 0.050 | 0.0375 | 0.0125 | 0.0303 | 0.603 |
